# Patient derived model of *UBA5-*associated encephalopathy identifies defects in neurodevelopment and highlights potential therapies

**DOI:** 10.1101/2024.01.25.577254

**Authors:** Helen Chen, Yong-Dong Wang, Aidan W. Blan, Edith P. Almanza-Fuerte, Emily S. Bonkowski, Richa Bajpai, Shondra M. Pruett-Miller, Heather C. Mefford

## Abstract

*UBA5* encodes for the E1 enzyme of the UFMylation cascade, which plays an essential role in ER homeostasis. The clinical phenotypes of UBA5-associated encephalopathy include developmental delays, epilepsy and intellectual disability. To date, there is no humanized neuronal model to study the cellular and molecular consequences of *UBA5* pathogenic variants. We developed and characterized patient-derived cortical organoid cultures and identified defects in GABAergic interneuron development. We demonstrated aberrant neuronal firing and microcephaly phenotypes in patient-derived organoids. Mechanistically, we show that ER homeostasis is perturbed along with exacerbated unfolded protein response pathway in cells and organoids expressing *UBA5* pathogenic variants. We also assessed two gene expression modalities that augmented *UBA5* expression to rescue aberrant molecular and cellular phenotypes. Our study provides a novel humanized model that allows further investigations of *UBA5* variants in the brain and highlights novel systemic approaches to alleviate cellular aberrations for this rare, developmental disorder.

**One-sentence summary:** Patient derived model of UBA5-assoicated DEE recapitulated disease phenotype, revealed defects in neurodevelopment, and highlighted potential therapies.

## Introduction

Mutations in the ubiquitin like modifier activating enzyme 5 (*UBA5*) gene have been associated with developmental and epileptic encephalopathy 44 (DEE44, OMIM: 617132), an autosomal recessive neurodevelopmental disorder characterized by severe developmental delay, hypotonia, spasticity, microcephaly, growth failure, and epilepsy (*1–11*). Most affected individuals achieve few developmental milestones and have intractable, early-onset seizures including infantile spasms, myoclonic, and motor seizures that can be life limiting. *UBA5* encodes a ubiquitously expressed E1 activating enzyme, first of a series of enzymes that is required for a process called ufmylation, a ubiquitin-like post-translational modification of proteins that results in the addition of ubiquitin fold modifier 1 (UFM1) (*12–14*). The ufmylation cascade involves three sequential steps to activate, conjugate, and ligate UFM1 onto substrates. First, proUFM1 is cleaved by a UFM1 specific peptidase 1/2 (UFSP1/2) to expose a C-terminal Gly; UBA5 activates UFM1 through an ATP-dependent adenylation of the newly exposed C-terminal Gly, forming a thioester intermediate (*12*). Next, UBA5 facilitates the conjugation of UFM1 onto the catalytic cysteine of the E2 UFM1-conjugase 1 (UFC1) through a transthiolation reaction (*12*). Lastly, the E3 UFM1-ligase 1 (UFL1) functions as a scaffold to bring UFC1-UFM1 conjugate to substrates that results in conjugation of UFM1 onto Lys residues on substrates (*13*). Much like ubiquitination, UFM1 modifications are reversible and can influence target protein interactions, stability, localization, and function (*14, 15*). Unlike ubiquitination, only one enzyme for each step of this cascade is known so far, thus no compensatory mechanisms are present (*12, 13*). Ufmylation has been implicated in a number of cellular processes including genome stability, vesicle trafficking, cell cycle progression, gene expression, and erythroid differentiation (*16–21*). The most prominent role for the ufmylation cascade is regulation of endoplasmic reticulum (ER) homeostasis and the unfolded protein response (UPR) pathway (*16, 21–26*). Conditional knockout of the ufmylation pathway components in mouse results in elevated ER stress and cell death (*8, 26*).

Complete lack of UBA5 activity is embryonic lethal in mice (*27*) and is likely lethal in humans. Recently, biallelic pathogenic variants in *UFSP2*, *UFM1* and *UFC1* have also been reported in individuals with microcephaly, global delays and seizures (*28, 29*). The ufmylation pathway is ubiquitous to all tissue, yet it is unclear why the major defects impact the central nervous system. To date, there are 38 individuals reported with *UBA5*-associated DEE (*1–11*). In nearly all cases, one variant is missense, resulting in a hypomorphic gene product, while the second variant is more deleterious, often causing premature truncation and resulting in nonsense mediated decay or producing a nearly non-functional gene product. Interestingly, over 65% of patients share the same variant for one of their two causative alleles: the missense variant, p.A371T, produces a hypomorphic protein that retains some normal function (*5, 7, 8*) (Fig. 1A). The p.A371T variant is present in the general population at a frequency of 0.27%, with a maximum allele frequency of 0.54% in the European Finnish population (Genome Aggregation Database, gnomAD v4.0 (*30*)). Furthermore, seven healthy adults with homozygous p.A371T variants have been reported; therefore, despite the decreased activity of the p.A371T variant, the combined activity of two p.A371T variants apparently provides enough activity to allow for normal cellular functions and prevents disease manifestation. This suggests that ‘boosting’ the amount of the p.A371T hypomorphic allele in affected individuals may be an effective therapy.

**Figure 1.**
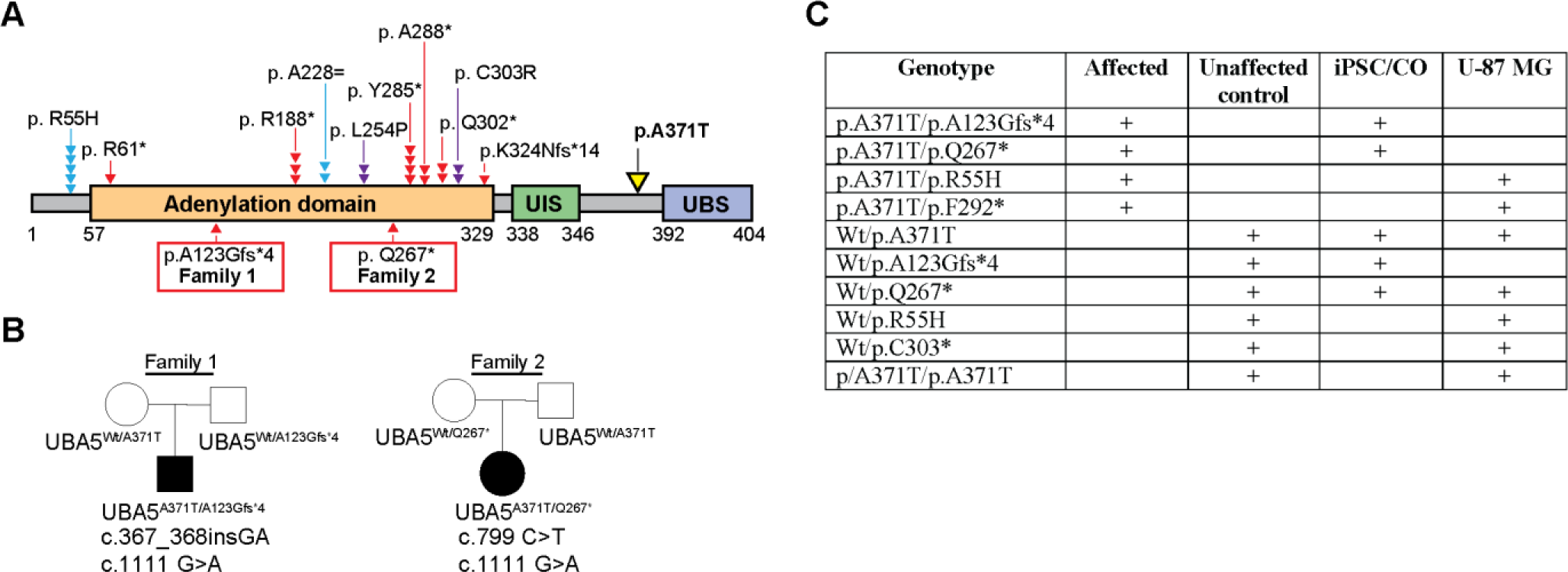
*UBA5* pathogenic variants and cell models included in this study. **(A)** Schematic representation of UBA5 structure with functional domains, UIS (UFM1-interacting motif); UBS (UFC1-binding sequence). All *UBA5*-DEE patients sharing the p.A371T variant (yellow) are listed. The second allele being either premature truncation (red) or splicing defect variants (blue) predicted to undergo nonsense mediated decay or missense (purple). Each red, blue or purple symbol represents one patient. **(B)** Family pedigree of probands included in this study. Circle: female; square: male; filled: affected proband. **(C)** Genotypes of *UBA5* cell models used in this study; all iPSC/CO are derived from patient or parent fibroblasts.

There have been major developments in therapies that modulate gene expression, with intervention at the gene, mRNA and protein stage. An important consideration in synthetic restoration of gene expression is ensuring that the level of upregulation is physiologically relevant. SINEUPs (Short Interspersed Nuclear Element UP regulation) are naturally occurring long noncoding antisense RNA molecules that modestly increase translation of overlapping mRNAs (*31–33*). Synthetic SINEUPs have been used to rescue motor deficits in a mouse model of GDNF-related Parkinson disease (*34*) and expression of FXN in a cellular model of Friedreich’s Ataxia (*35*). CRISPR (clustered regularly interspaced short palindromic repeat) and CRISPR-associated (Cas) protein system has been adapted for human genome editing (*36*). Combining single guide RNA (sgRNA) along with a nuclease-null Cas9 (dCas9) can up- or down-modulate transcription of target genes without inducing a double-stranded break (*37*). The CRISPRa-dCas9 system has been utilized to rescue haploinsufficient obesity disorder (*38*) and Smith-Magenis syndrome, a severe neurodevelopmental disorder (*39*), in mice.

Due to the limited number of patients and the inaccessibility of affected brain tissue, investigations of *UBA5*-associated DEE have thus far only been performed using non-mammalian models (*7, 11, 40*) or non-neuronal human tissue (*6–8*). Thus, there is a gap in knowledge regarding how *UBA5* pathogenic variants affect human neurodevelopment. More importantly, there is no systemic treatment available for patients, only selected drugs to combat symptoms. In this study, we identified two previously unpublished probands with biallelic *UBA5* pathogenic variants and generated 3D cortical organoids (CO) from proband-derived induced pluripotent stem cells (iPSC) along with healthy parental controls to investigate molecular and functional alterations during cortical development. Our findings demonstrate that *UBA5* proband COs are smaller in size and exhibit aberrant neuronal firing phenotypes in these *in vitro* humanized models. We identify a significant reduction of γ-aminobutyric acid (GABA)ergic interneurons in *UBA5* proband CO. Along with proband-derived CO, we also engineered U-87 MG cell lines that represent *UBA5* patient variants and show exacerbation of the UPR pathway and perturbation of ER homeostasis. Given the hypothesis that the p.A371T variant does not cause DEE in the homozygous state (*5*), we generated a cell line homozygous for the p.A371T variant and show that it behave as wildtype cells. As a proof of concept, we utilized two approaches, SINEUP and CRISPRa, to moderately increase the expression of p.A371T, and we were able to ameliorate defects associated with ER homeostasis and neuronal firing in *UBA5* proband CO models. Collectively, our study provides a novel reliable human model to investigate *UBA5*-associated DEE, recapitulating hallmarks of the disease while providing new insights into the molecular mechanisms of this ultrarare disorder. Most importantly, we introduce two novel therapeutic approaches that could potentially be used in treatment development for *UBA5*-associated DEE.

## RESULTS

### Identification of two patients with compound heterozygous pathogenic variants in the *UBA5* gene

We report two previously unpublished affected individuals with biallelic pathogenic variants in *UBA5* (Fig. 1A and 1B). Individual 1 is a 6-year-old with intractable epilepsy, severe developmental delay, spasticity, dystonia, and cortical visual impairment. The individual was noted to have early delays with feeding difficulties shortly after birth, poor head control, hypotonia, and decreased engagement in the first few months after birth. The individual developed seizures at 3 months that have been intractable to multiple antiseizure medications and vagal nerve stimulation. Seizures include ‘jack knife’ seizure with flexion and stiffening of his body, startle seizures, and tonic seizures. Initial EEG showed hypsarrhythmia; most recent EEG shows multifocal epileptiform discharges, paroxysmal fast activity, and abnormal background with generalized slowing. MRI shows thin corpus callosum and delayed myelination. Head circumference at 4 and 5 years is at ∼25^th^ percentile. The individual has excessive drooling treated with botulinum toxin injections and clonus treated with baclofen. Feeding is by gastrostomy-jejunostomy tube and BiPAP is used at night for obstructive sleep apnea. Developmentally, the individual is severely delayed: makes sounds but is nonverbal; uses eye gaze for “yes” and “no” communication; and is nonambulatory but uses a stander with assistance. Trio exome sequencing revealed biallelic variants in the *UBA5* gene (transcript NM_024818.3): c.367_368insGA, p.(A123Gfs*4) / c.1111G>A, p.(A71T).

Individual 2 is a 9-year-old with intractable epilepsy, profound developmental delay, cortical visual impairment, and spastic quadriplegia. This individual developed infantile spasms at 4 months of age, later developing tonic seizures, myoclonic jerks, and atypical absence seizures, all of which have been refractory to antiseizure medications and the ketogenic diet. EEG shows slow spike and wave discharges with increase in slow-wave sleep; she exhibits paroxysmal fast activity with tonic seizures. Brain MRI shows thinning of the corpus callosum and findings consistent with slowly progressive cerebral volume loss. The individual has microcephaly, with head circumference below 3^rd^ percentile since at least 6 months old. Developmentally, the individual is profoundly delayed: has minimal interaction, is nonverbal and unable to sit, stand or walk independently; is fed by gastrostomy tube. Spastic quadriplegia is treated with botulinum toxin injections to pectoralis major and hip adductors bilaterally. Trio exome sequencing revealed biallelic variants in the *UBA5* gene (transcript NM_024818.3): c.799C>T, p.(G267*) / c.1111G>A, p.(A371T).

### Modelling *UBA5* DEE using patient-derived cortical organoids shows deficit in GABAergic pathway processes

Previous investigations of *UBA5* pathogenic variants utilized patient-derived fibroblasts (*7, 8*), HEK293 cells (*7*), and *Drosophila* (*7, 8, 40*) to model various aspects of the disorder. However, these models are not able to recapitulate human neurodevelopment. To establish a model that is suitable to study the impact on human neurodevelopment, we generated cortical organoids (CO) from patient-derived iPSC by reprogramming fibroblasts from the UBA5^A371T/A123Gfs*4^ and UBA5^A371T/Q267*^ patients and their healthy parents (as controls) (Fig. 1C). iPSC lines were assessed for the expression and localization of key pluripotency markers, and chromosomal abnormalities (Fig. S1).

We first confirmed decreased expression of *UBA5* mRNA in proband CO. We observed no changes in *UFM1, UFC1* and *UFBP1*, which are members in the ufmylation pathway (Fig. S2A), consistent with previous investigation of fibroblasts from other *UBA5* patients (*8*). Next, we confirmed expressions of cortical markers, including *FEZF2, BRN2, DCX, SATB2* and *CTIP2* (Fig. S2B). To interrogate potential defects in corticogenesis, we performed the Gene Set Enrichment Analysis (GSEA), comparing bulk RNAseq findings from proband and control (unaffected parents) CO. Pathway analysis revealed that GABA neurotransmission and receptor signaling were among the top processes that were significantly downregulated in both proband CO compared to controls (Fig. 2A), and the genes involved in GABA receptor signaling were significantly downregulated in proband CO in both families (Fig. 2B). In UBA5^A371T/Q267*^ proband CO (Family 2), we also noted a significant increase in processes related to ER homeostasis and UPR (Fig. 2A).

**Figure 2.**
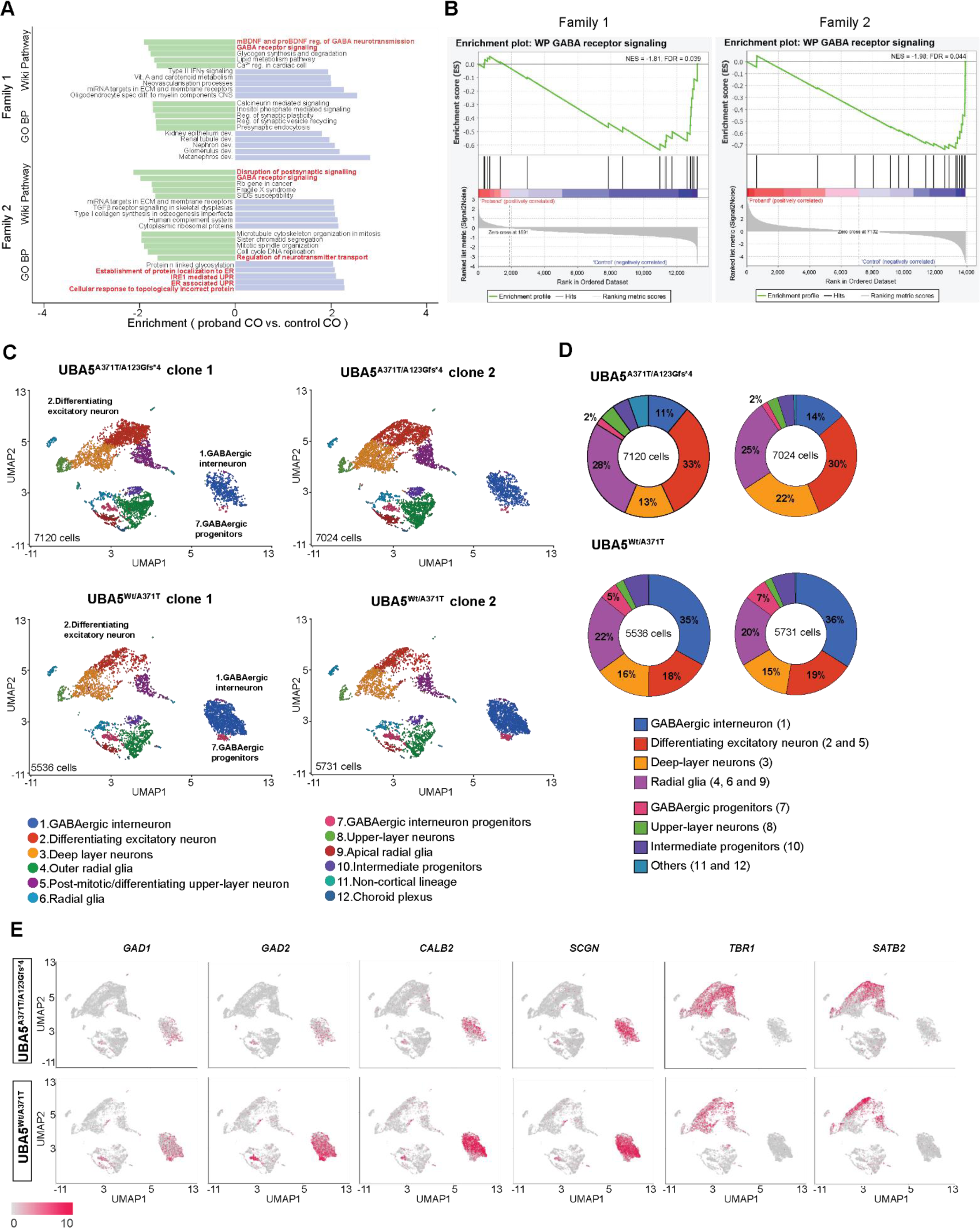
Bulk and single-cell RNA sequencing reveal a deficit in GABAergic interneurons processes in 100-day old CO derived from *UBA5* proband. **(A)** Normalized enrichment scores of gene sets between proband and control CO from families 1 (UBA5^A371T/A123Gfs*4^) and 2 (UBA5^A371T/Q267*^) as determined by GSEA. The top five pathways from GO Biological Process and Wiki pathways databases are shown. **(B)** The enrichment plots for GABA receptor signaling from Wiki pathways in families 1 (UBA5^A371T/A123Gfs*4^) and 2 (UBA5^A371T/Q267*^). **(C)** UMAP plot showing unbiased clustering of cell types, 31,950 cells. **(D)** Organoid cell type contribution for between genotypes, cluster identify from **(C)** as indicated. **(E)** Feature plots showing reduced expression of GABAergic interneuron genes in *UBA5* proband compared to control CO (both clones are combined in each plot).

### Single-cell profiling of *UBA5* proband-derived CO shows deficit in GABAergic neuron population

To better understand potential defects in cortical development, we performed single-cell RNA sequencing (scRNAseq) to assess cellular diversity and cell-type specific transcriptomic signatures in day 100 CO from Family 1. We included 2 different iPSC clones from UBA5^A371T/A123Gfs*4^ proband and UBA5^Wt/A371T^ control. Using unsupervised clustering, we identified 12 composite clusters using canonical marker genes (Fig. 2C, Table S1). We identified two clusters of inhibitory GABAergic interneurons of the ganglionic eminence and four clusters of excitatory neurons with identity matching specific cortical layers. We identified three clusters of radial glial population, with a predominant late-stage, outer radial glial population (cluster 4) expressing *TNC, FAM107A* and *HOPX*; early-stage radial glial population (cluster 6) expressing *TAGLN2* and *CRYAB,* and apical radial glia population expressing *BHLHE40* and *PLOD2* (cluster 9) (*41*). Astrocytes are present in cluster 6 as identified by *S100B* and *GFAP*. Intermediate progenitors (cluster 10) were identified based on expressions of *NEUROG1, EOMES* and *MKI67* (*42*). We also detected a small cluster of cells (<1%) with choroid plexus identity (cluster 12) that expressed *MGST1* and *FIBIN*(*43*). Cells within cluster 11 (<1%) expressed markers indicating cerebellum/rostral hindbrain identity, such as *PAX2* and *BARHL* (*44, 45*).

Most strikingly, we observed a drastic reduction in the relative proportion of GABAergic interneurons of the ganglionic eminence in *UBA5* proband CO (∼15%) compared to healthy parental controls (∼40%) (Fig. 2C and 2D). We identified two clusters of GABAergic interneurons expressing *GAD1, GAD2, CALB2, DLX5, DLX2* and lacking *NEUROD2* and *NEUROD6* (*46–49*). These two clusters of GABAergic interneurons are classified as early progenitors expressing *GAD2, NPY* and *GSX2* (cluster 7) and mature interneurons expressing *CALB2* and *SCGN* (cluster 1). Expression analysis of GABAergic markers showed reduction in proband CO, with *GAD2*, *SCGN* and *CALB2* being most drastically decreased (Fig. 2E and Fig. S4). On the other hand, we identified subtypes of glutamatergic excitatory neurons residing in specific cortical layers, which are significantly increased in proband CO compared to control CO (Fig. 2C and 2D). For example, cluster 2 represents differentiating excitatory neurons as identified by *MEF2C, TBR1* and *SATB2* and constituted ∼23% of cells within proband CO, but only 12% in control CO. The proportion of *TBR1* and *NRN1*-positive deep layer excitatory neurons in cluster 3 was also increased from 15% in control CO to 22% in proband CO. We did not observe any differences in cluster 5 and 8, which were differentiating upper-layer excitatory neuron and upper-layer excitatory neurons, respectively, and together constituted ∼10% of cells. Collectively, these findings further highlight aberrant neocortex development due to *UBA5* pathogenic variants. Lastly, we noted no difference in the proportion of intermediate progenitors or radial glial cells between proband and control CO (Fig. 2C and 2D). Our findings were reproducible across both clones in proband and healthy parental control.

### GABAergic markers are reduced in D100 CO derived from *UBA5* probands

To validate whether expression of GABAergic and glutamatergic markers are altered in CO derived from *UBA5* probands, we performed qRT-PCR and immunoblotting analyses on 100-day old CO. Immunoblotting identified two isoforms of UBA5: a ∼45kDa protein (NP_079094.1) encoded by a 4692-nt canonical transcript (NM_024818.6), and a ∼39kDa protein (NP_938143.1) encoded by a 4563-nt transcript (NM_198329.4), which is the minor contributor. Both isoforms of UBA5 are significantly reduced in UBA5^A371T/A123Gfs*4^ and UBA5^A371T/Q267*^ proband CO compared to UBA5^Wt/A371T^ CO (Fig. 3C). In the two other healthy parental control CO, UBA5 expression is also reduced to half, likely due to nonsense mediated decay of the mutant allele. In proband CO, qRT-PCR analysis revealed significant decreased levels of GABAergic markers, *GAT1, GAD2*, *GAD1*, *GABRA1* and *SLC32A1* (Fig. 3A), findings supported by orthogonal immunoblotting assays (Fig. 3B and Fig. S2C). Analysis of glutamatergic markers showed no differences between proband and control CO. Thus, the observed increase in glutamatergic excitatory neurons in our scRNAseq analysis is likely proportional, not absolute change, whereas the total number of GABAergic interneurons are reduced in proband CO. Next, we performed whole mount immunofluorescent staining of SATB2 and CTIP2 after tissue clearing to assess CO structures (Fig. 3C). Quantification of layer-specific neurons showed the proportion of both SATB2+ and CTIP2+ cells are similar between proband and healthy control CO (Fig. 3D)

**Figure 3.**
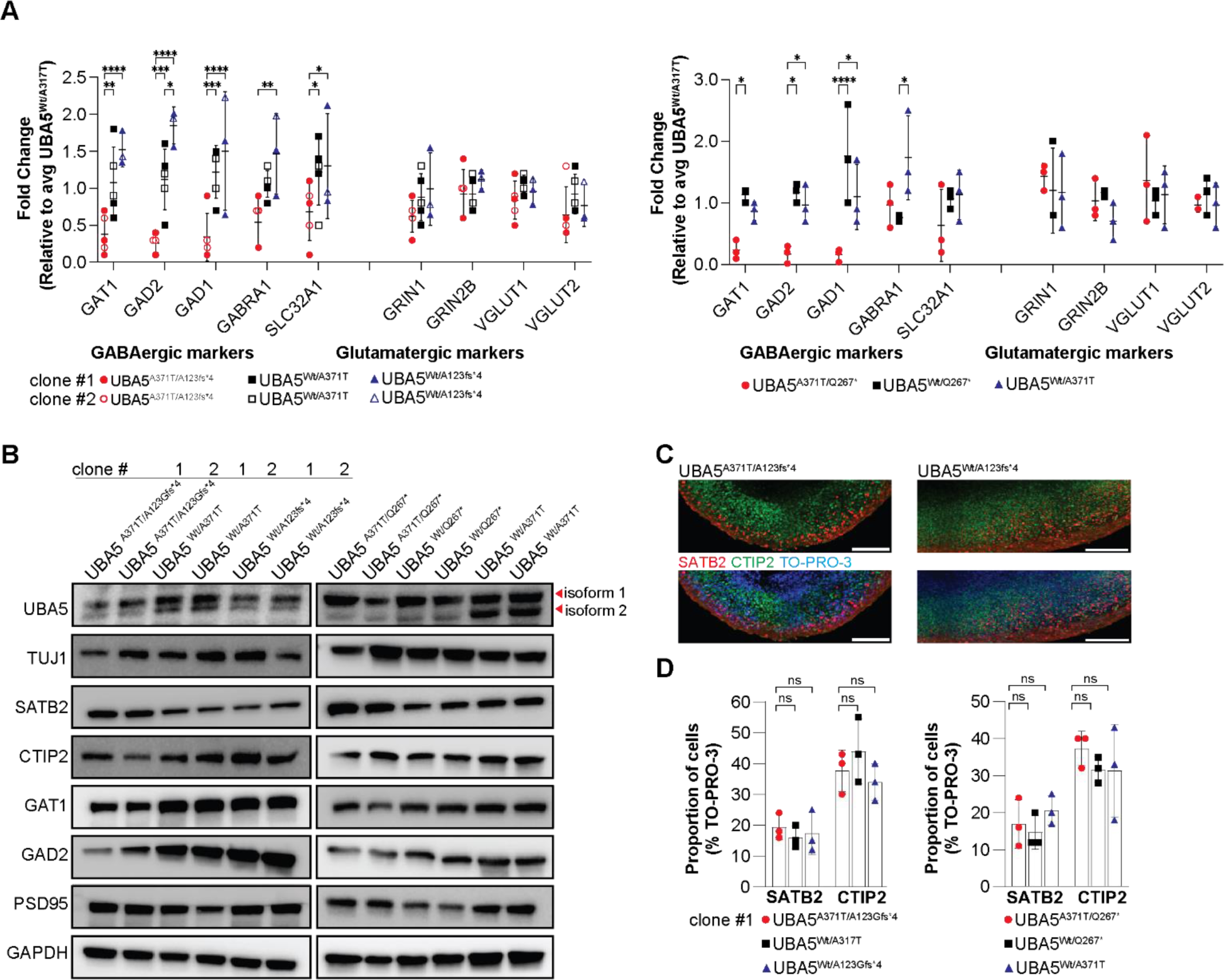
GABAergic markers are reduced in 100-day old CO derived from *UBA5* probands. **(A)** Transcript level of GABAergic markers are significantly reduced in CO derived from *UBA5* probands compared to control CO, normalized to UBA5^Wt/A371T^. Each data point represents three CO, plotted as mean ± SD. *P<0.05, **P<0.01, ***P<0.001 and ****P<0.0001. **(B)** Immunoblot analysis shows reduced expression of both isoforms of UBA5 and confirms reduced expression of GAD2 in CO derived from *UBA5* probands compared to control CO. GAPDH served as loading control. All images shown in the manuscript are representative images of 3 independent experiments. **(C)** Staining for cortical layer identities of CO shows later-born surface-layer neurons (SATB2) populate the superficial regions of the organoid, whereas early-born deep-layer neurons (CTIP2) populate the inner regions of the organoid. Scale bars: 50 µm. **(D)** Proportion of cells expressing SATB2 or CTIP2 in CO. Each data point represents one CO, plotted as mean ± SD. ns: not significant.

### CO derived from *UBA5* probands showed reduced growth and aberrant neuronal activity

To investigate whether proband CO modelled progressive microcephaly as seen in *UBA5* patients (*7, 8*), we measured the diameter of CO from day 50 to 120 (Fig. 4A). Proband CO was consistently 25% smaller than control CO. Between day 50 and 120, proband and control CO all grew by ∼40%, with no differences in the growth rate. Using our differentiation protocol, we demonstrated that the proband CO recapitulates morphological features of microcephaly seen in *UBA5* patients (*4, 5, 7, 8*).

**Figure 4.**
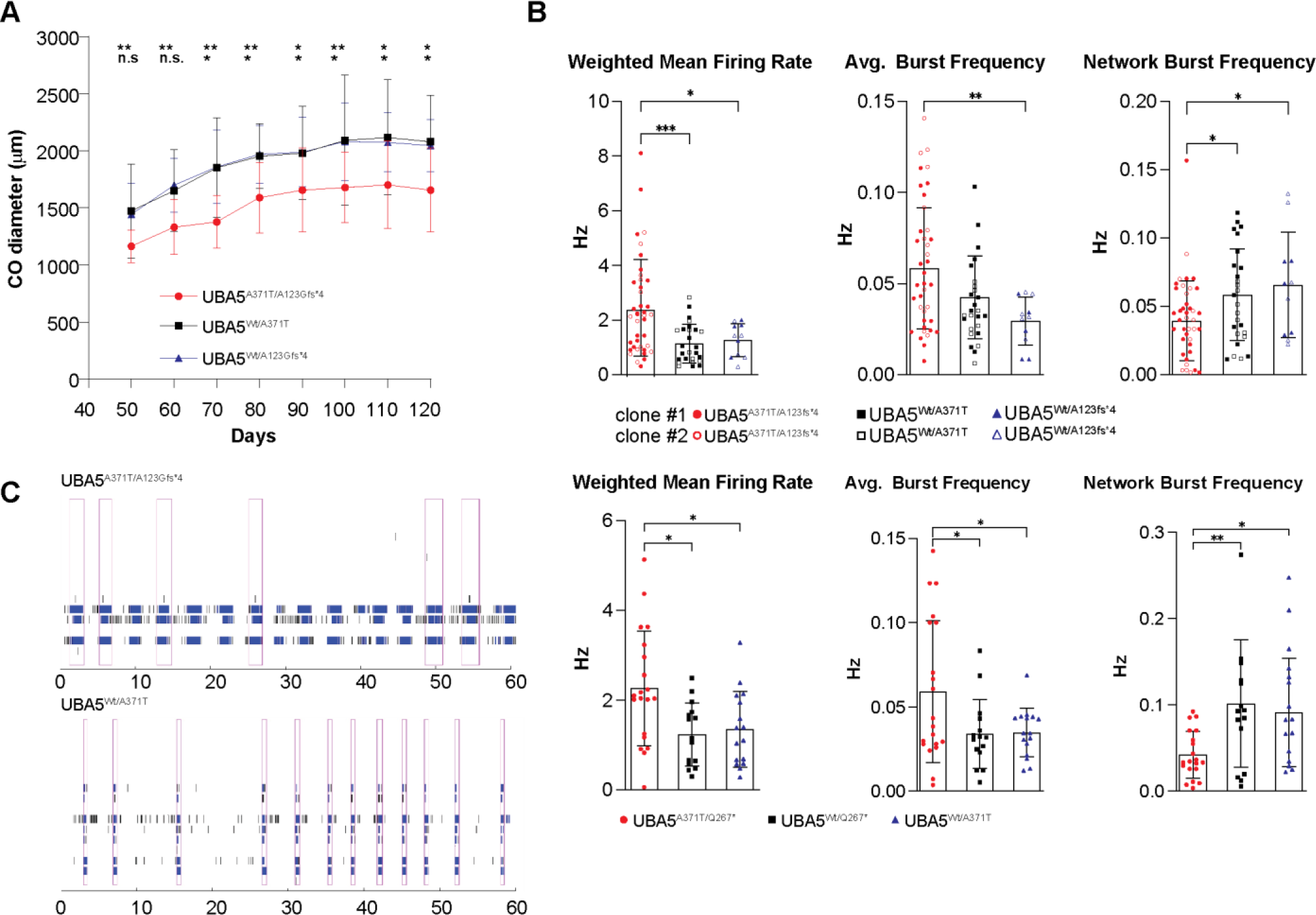
CO derived from *UBA5* proband showed reduced size and aberrant network activity. **(A)** Growth trajectory of CO as measured by CO diameter. The same 11-15 CO were measured from D50 to 120, plotted as mean ± SD. *P<0.05 and **P<0.01. ns: not significant. **(B)** Functional characterization of D110 CO derived from *UBA5* probands using MEA showing changes in weighted mean firing rate, averaged burst frequency and network burst frequency compared to control CO. Each data point represents one CO, plotted as mean ± SD. *P<0.05, **P<0.01 and ***P <0.001. **(C)** Representative raster plot of neural network activity. Each black tick indicates the time a spike occurred; blue tick indicates the spikes are part of a burst and magenta rectangles indicate a network burst event. Each row represents a recording from an electrode.

One clinical feature of *UBA5* associated DEE is abnormal EEG and seizures (*5, 7, 8*). To assess spontaneous neuronal activity, we performed recordings of CO using a multi-electrode array (MEA) (Fig. 4B). On day 100, one CO was plated per well and recordings were performed beginning on day 110. Proband CO exhibited consistent increases in weighted mean firing rate and averaged burst frequency. Interestingly, we noted proband CO exhibited reduced network burst frequency (Fig. 4C). Network behavior is an important aspect of neurodevelopment (*50–52*), and epileptic seizure is not a synchronous hyperactive state (*53–55*). We performed recording of the same CO every 10 days for a 35-day period and observed persistent aberrant neuronal activity (Fig. S5). Moreover, we noted these neuronal defects in both probands. Thus, increased neuronal firing and burst frequency, along with decreased network bursts, are shared features in *UBA5* cellular models.

### *UBA5* pathogenic variants disrupt the ufmylation pathway in engineered U-87 MG cells

To investigate the molecular aberrations of *UBA5* pathogenic variants, we generated U-87 MG cells carrying compound heterozygous pathogenic variants in *UBA5* (Fig. 1C). We generated two cell lines, UBA5^A371T/R55H^ and UBA5^A371T/F292*^, to represent p.A371T/loss-of-function genotypes, along with heterozygous control lines (UBA5^Wt/A371T^, UBA5^Wt/R55H^ and UBA5^Wt/C303*^). The UBA5^A371T/R55H^ variant was previously identified in five individuals in two families; the R55H missense variant produced by the c.164G>A point mutation alters splicing and causes exon skipping of exon 2, leading to nonsense-mediated decay (*8, 56*). UBA5^A371T/F292*^ was generated to model compound heterozygous p.A371T with a premature truncation variant, present in 12 individuals (*3, 6–8*). Most importantly, we generated a UBA5^A371T/A371T^ line to compare homozygous p.A371T variant with wildtype *UBA5*, given that individuals homozygous for p.A371T do not present features associated with *UBA5*-DEE (*5, 7, 8*). Immunoblotting analysis detected reduced abundance of both isoforms of UBA5 in UBA5^A371T/R55H^, UBA5^A371T/F292*^, UBA5^Wt/R55H^, and UBA5^Wt/C303*^ compared to UBA5^Wt/Wt^ cells as predicted (Fig. 5A and Fig. S6C). Transcript level and protein abundance of other components of the ufmylation pathway were not altered in UBA5^A371T/R55H^ and UBA5^A371T/F292*^ cells (Fig. 5A and Fig. S6A). Immunofluorescence staining showed similar cytoplasmic localization of UBA5 in all cell lines (Fig. S6B). To determine the functional capacity of the E1-like activity of UBA5 in cells, we performed immunoblotting assays with both reduced and non-reduced lysates to detect UFM1-UBA5 and UFM1-UFC1 conjugates (Fig. 5B). Together, these findings demonstrate that cells expressing compound heterozygous *UBA5* mutations exhibit significant reduction of ufmylation activity, while homozygous UBA5^A371T/A371T^ cells showed slight reduction, though not significant, further highlighting the milder effects of the p.A371T variant.

**Figure 5.**
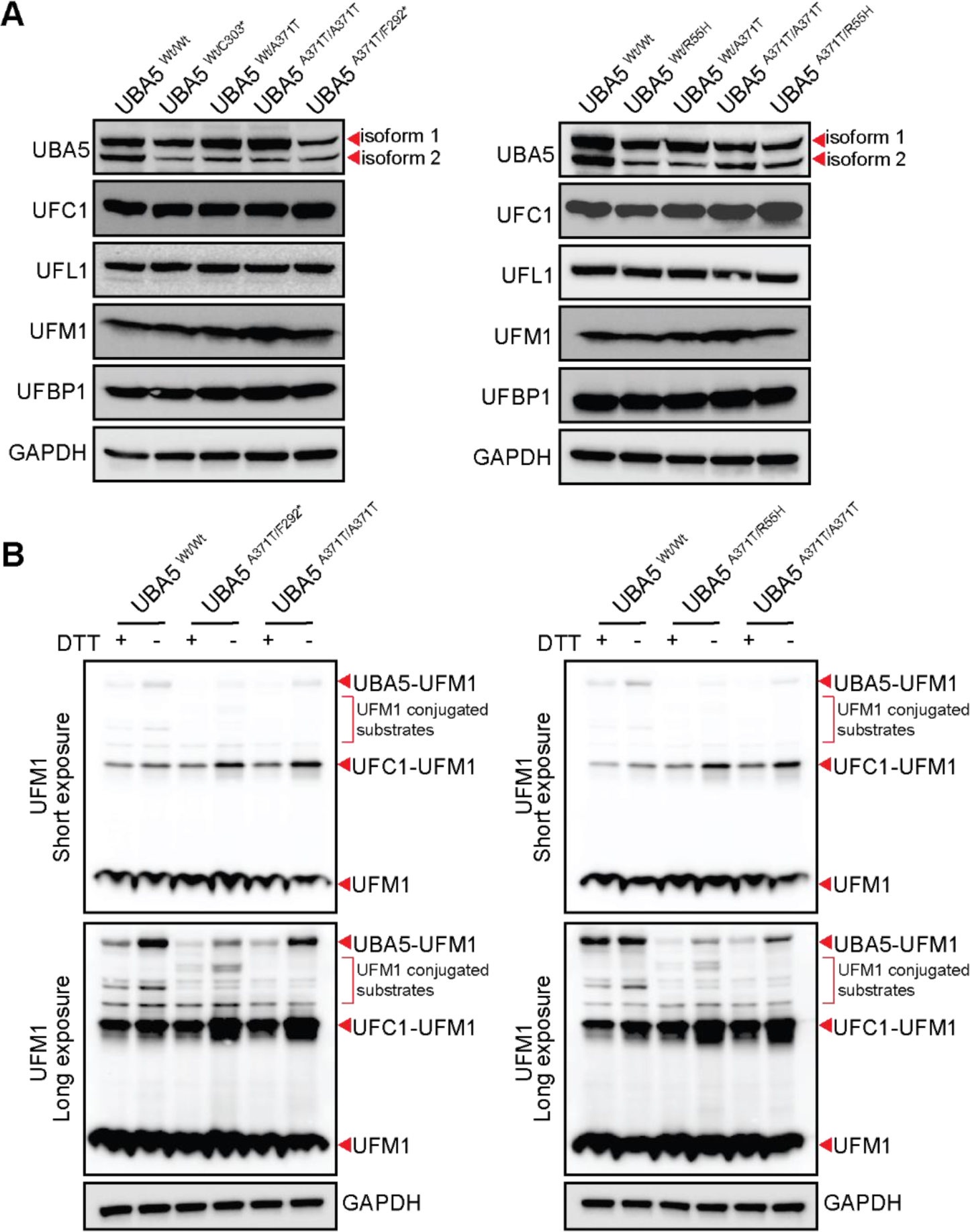
Characterization of the ufmylation pathway in U-87 MG cells expressing *UBA5* pathogenic variants. **(A)** Immunoblot analysis of various components of the ufmylation pathway, showing decreased UBA5 levels for both isoforms in cells expressing UBA5^A371T/F292*^ and UBA5^A371T/R55H^, with no changes observed in cells expressing UBA5^A371T/A371T^ compared to wildtype. GAPDH served as loading control. All images shown in the manuscript are representative images of 3 independent experiments. **(B)** Immunoblot analysis of UFM1 conjugates with and without reducing agent, DTT. GAPDH served as loading control. All images shown in the manuscript are representative images of 4 independent experiments.

### *UBA5* pathogenic variants disturb ER homeostasis in U-87 MG cells and CO

ER homeostasis is essential to the maintenance of intracellular calcium and the biosynthesis, folding and transport of proteins, lipids and sterols (*57*). Various cellular aberrations can lead to ER stress and trigger the unfolded protein response (UPR) to alter transcriptional and translational programs in cells (*58*). However, if UPR fails to restore ER homeostasis, apoptotic programs are then activated as a result (*58*). UPR relays four interlinked mechanisms to accommodate protein folding during ER stress: 1) translation attenuation (*59, 60*); 2) increase expression of ER chaperones (*60*); 3) induction of ER-associated degradation (ERAD) machinery (*61*) and 4) ER size expansion. The ufmylation pathway is implicated in the maintenance of ER homeostasis and stress response (*16, 21–26*). To better understand the cellular consequences of *UBA5* variants, we examined signaling pathways dictated by the three main proteins that relay UPR at the ER membrane: 1) double-stranded RNA-activated protein kinase (PKR)-like endoplasmic reticulum kinase (PERK); 2) activating transcription factor 6 (ATF6) and 3) inositol requiring kinase 1 (IRE1). PERK phosphorylates ribosomal initiating factor (eIF2α), and p-eIF2α will attenuate protein translation and promote the translation of activating transcription factor 4 (ATF4), which promotes the expression of ER chaperones, ERAD proteins and apoptosis factors (*59, 60*). We observed a significant increase in the active form of PERK (p-PERK) and p-eIF2α in cells expressing *UBA5* pathogenic variants, UBA5^A371T/R55H^ and UBA5^A371T/F292*^(Fig. 6A and Fig. S7A). The second effector is ATF6, which translocates to the nucleus to elicit transcription upregulation of UPR target genes that overlap with those activated by ATF4 (*61–63*). There is increased nuclear localization of ATF6 in UBA5^A371T/R55H^ and UBA5^A371T/F292*^ cells (Fig. 6B), along with increased nuclear expression of a downstream target, CHOP (Fig. 6C). We observed augmented expression of other target genes such as BiP and GRP94 (Fig. 6A and Fig. S7A). The last main branch of UPR is via IRE1, which autophosphorylates to activate its endoribonuclease activity, and then splices *XBP1* mRNA to produce a transcription factor that promotes gene expression of UPR target genes (*64*). In UBA5^A371T/R55H^ and UBA5^A371T/F292*^ cells, we noted significant decrease in the abundance of both IRE1α and phosphorylated IRE1α (p-IRE1α) (Fig. 6A and Fig. S7A), consistent with previous studies showing IRE1α stability is dependent on the ufmylation pathway (*65, 66*). We quantified that the amount of spliced *XBP1* is significantly less in both UBA5^A371T/R55H^ and UBA5^A371T/F292*^ cells (Fig. S7B), consistent with the notion that the endoribonuclease activity of IRE1α is reduced. Lastly, we measured ER size using calnexin (Fig. 6D and 6E), a membrane chaperon protein whose abundance was not altered by *UBA5* pathogenic variants (Fig. 6A and 6D) and noted a significant increase in relative ER size in UBA5^A371T/R55H^ and UBA5^A371T/F292*^ cells. Expansion of ER volume was previously observed in fibroblasts from *UBA5* patients (*7*). Prolonged, unresolved ER stress results in activation of the caspase cascade and then apoptosis (*67*); immunoblotting analysis showed increase in the cleavage of poly(ADP)-ribose polymerase (PARP) in UBA5^A371T/R55H^ and UBA5^A371T/F292*^ lysates (Fig. 6A and Fig. S7A), suggesting that *UBA5* pathogenic variants cause apoptosis due to unresolved ER stress. Examination of UBA5^A371T/A371T^ cells showed no differences in the expression of UPR proteins and ER swelling (Fig. 6A-E and Fig. S7A) when compared to wildtype cells, further contributing to the notion that homozygous expression of the p.A371T is not sufficient to cause cellular defects. Furthermore, we observed a similar increase in p-PERK and p-eIF2α levels and decrease in IRE1α level in day-100 proband CO compared to control CO (Fig. 6F and Fig. S7C). Together, these findings suggest that ER homeostasis is perturbed with exacerbated UPR in cells expressing pathogenic *UBA5* variants.

**Figure 6.**
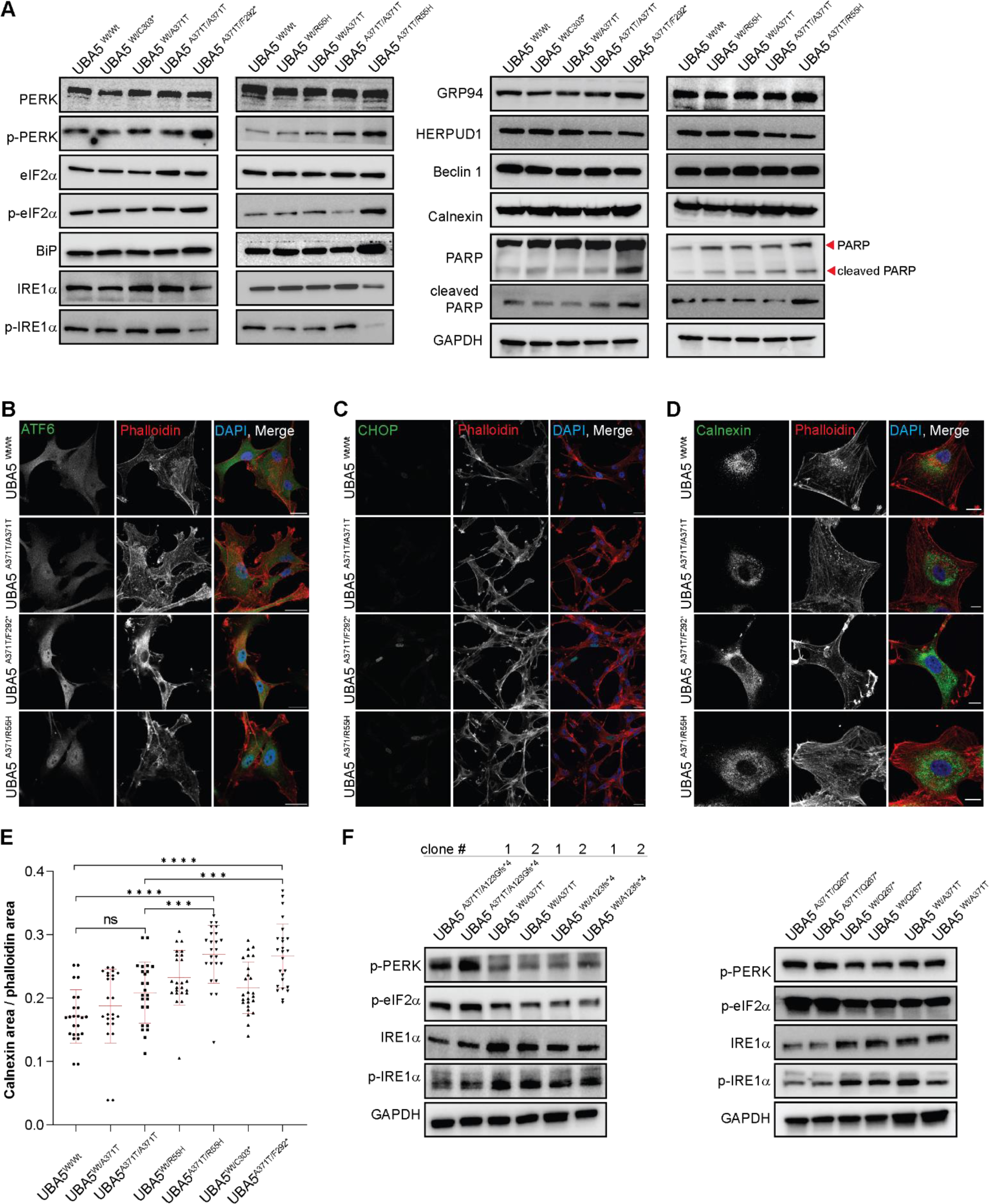
U-87 MG cells expressing *UBA5* pathogenic variants exhibit perturbed ER homeostasis. **(A)** Immunoblot analysis showing elevated expression of various components of the UPR pathway in cells expressing UBA5^A371T/F292*^ and UBA5^A371T/R55H^, with no changes observed in cells expressing UBA5^A371T/A371T^ compared to wildtype. GAPDH served as loading control. All images shown in the manuscript are representative images of 3 independent experiments. **(B-D)** Representative images of immunofluorescent staining with **(B)** ATF6 and phalloidin, **(C)** CHOP and phalloidin and **(D)** calnexin and phalloidin in U-87 MG cells. Scale bars: 10 µm. **(E)** Quantitation of relative ER area as measured by calnexin area/phalloidin area from (**D**). Each data point represents one image, plotted as mean ± SD. ***P<0.001, ****P<0.0001 and ns: not significant. **(F)** Immunoblot analysis showing elevated expression of various components of the UPR pathway in 100-day old proband CO. GAPDH served as loading control. All images shown in the manuscript are representative images of 3 independent experiments.

### Pathogenic *UBA5* variants respond to small molecule-induced ER stress similar to wildtype *UBA5*

Next, we sought to determine whether cells expressing *UBA5* pathogenic variants can recognize and respond to induced ER stress given that ER homeostasis is disturbed. We utilized two small molecules to induce two different sources of ER stress: Thapsigargin (TG) raises intracellular calcium while depleting ER calcium storage, and Bafilomycin A (BFA) causes cytoplasmic accumulation of H^+^ by inhibiting V-ATPase. We added either small molecule to induce ER stress in U-87 MG cells, then quantified key UPR markers by immunoblotting or immunofluorescence. We noted significant increase in BiP, HERPUD1, p-PERK and p-eIF2α upon TG or BFA stimulation in all cell lines examined (UBA5^Wt/Wt,^ UBA5^A371T/A371T^, UBA5^A371T/R55H^ and UBA5^A371T/F292*^; Fig. S8A). Immunofluorescence images show increased nuclear ATF6 and CHOP in all cell lines, suggesting that response to TG- or BFA-induced ER stress is similar regardless of *UBA5* genotype (Fig. S8B and S8C). Together, these findings show that despite perturbed ER homeostasis caused by *UBA5* pathogenic variants, the cells are still able to generate an appropriate response upon small molecule-induced ER stress.

### Application of synthetic SINEUP increases abundance of UBA5 gene product and restores ER homeostasis in U-87 MG cells

We designed two synthetic SINEUP constructs that are specific for *UBA5* mRNA (isoform 1: NM_024818.6) based on a previously published design that targets *eGFP* mRNA (*33*). Both synthetic SINEUP constructs contains a 72-nucleotide 5’ pairing sequence as the binding domain that confers *UBA5* specificity, and a 167-nucleotide interspersed nuclear element (SINE) B2 sequence as the effector domain together on a lentivirus-ready plasmid (Fig. 7A). We introduced a SINEUP construct or a non-targeting control (NTC) to cells expressing UBA5^A371T/R55H^ and UBA5^A371T/F292*^ via lentiviral transduction and examined UBA5 expression after 72 hours (Fig. 7B). We noted a modest ∼1.5-fold increase in UBA5 expression with SINEUP treatment compared to NTC (Fig. 7C). UBA5 expression following SINEUP is similar to cells expressing UBA5^A371T/A371T^ (Fig. 7B), and response to SINEUP treatment is specific for isoform 1 (as designed). As expected, we see no changes to *UBA5* mRNA with SINEUP treatment, indicating a translational enhancement (Fig. S9A). We observed increased UBA5-UFM1 conjugates after SINEUP application (Fig. S9B). Additionally, we continue to see augmented UBA5 expression up to 12 days after application of SINEUP, but with a minimum 3.5-fold increase instead, likely due to multiple integration of the SINEUP constructs into the genome (Fig. S9C).

**Figure 7.**
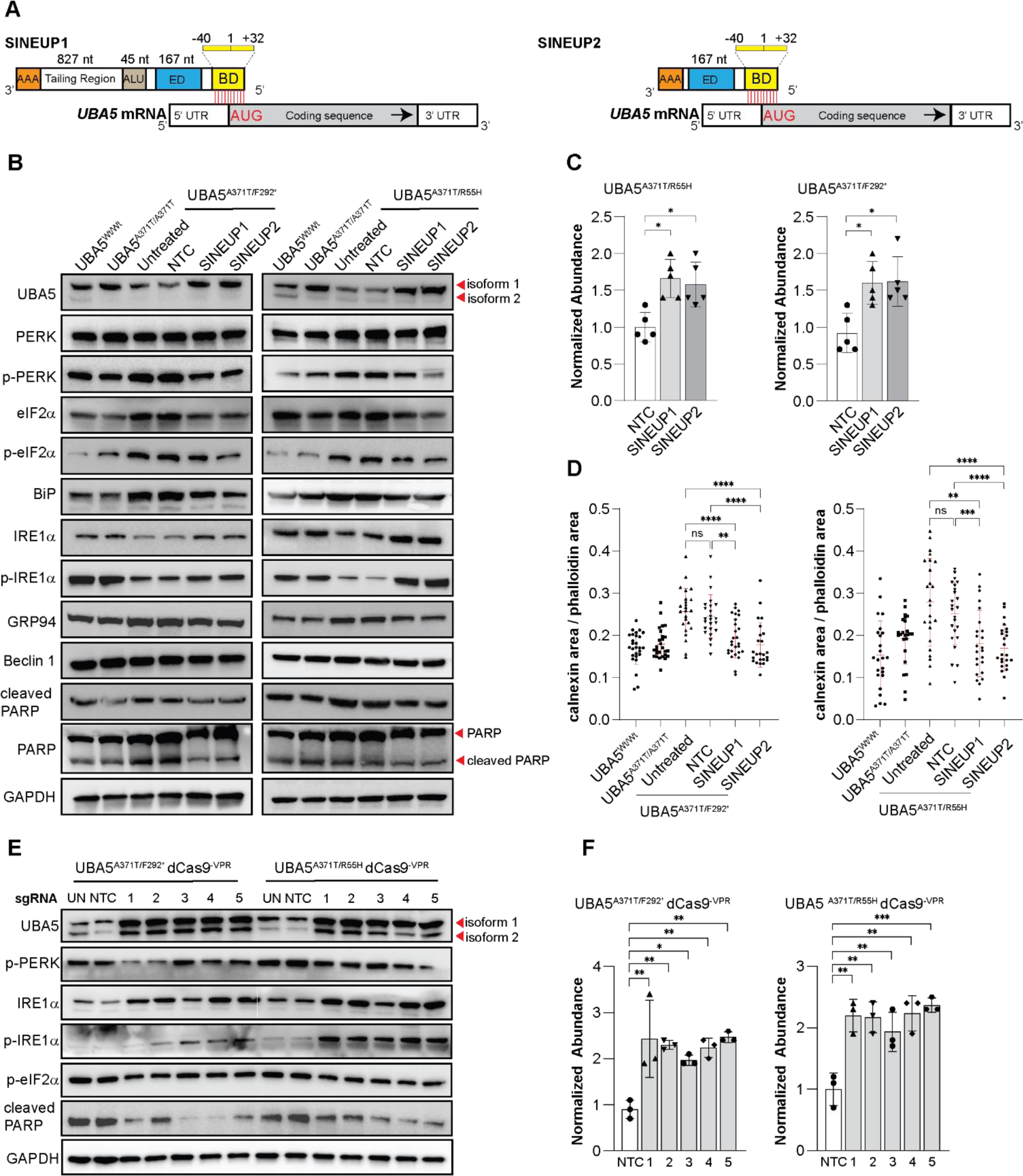
Genetic rescue of *UBA5* expression restores ER homeostasis in U-87 MG cells expressing *UBA5* pathogenic variants. (**A)** Schematic representation of synthetic SINEUP. SINEUP binding domain (yellow box: BD; 72nt) overlaps, in antisense orientation to sense coding sequence *UBA5* mRNA (grey box). SINEUP effector domain (blue box: ED; 167nt) contains an inverted SINBE2 element. Other components include partial ALU repeats (brown box: ALU; 45nt), tailing region (white box; 827nt) and a poly (A) tail (orange box: AAA). **(B)** Immunoblot analysis showed restored expression UBA5 and UPR pathway in cells following 72 hr of SINEUP treatment. NTC served as negative control for SINEUP. GAPDH served as loading control. All images shown in the manuscript are representative images of 3-5 independent experiments. **(C)** Quantification of UBA5 protein abundance following SINEUP treatment, first normalized to GAPDH, and then to averaged NTC treatment. Each data point represents one experiment, plotted as mean ± SD. *P<0.05. **(D)** Quantitation of relative ER area as measured by calnexin area/phalloidin area in cells following 72 hr of SINEUP treatment showed reduction in ER expansion. NTC served as negative control for SINEUP. Each data point represents one image, plotted as mean ± SD. **P<0.01, ***P<0.001, ****P<0.0001 and ns: not significant. **(E)** Immunoblot analysis showed restored expression UBA5 and UPR pathway in cells following 72 hr of sgRNA treatment. NTC served as negative control for sgRNA. GAPDH served as loading control. All images shown in the manuscript are representative images of 3 independent experiments. **(F)** Quantification of UBA5 protein abundance following sgRNA treatment, first normalized to GAPDH and then to NTC treatment. Each data point represents one experiment, plotted as mean ± SD. *P<0.05, **P<0.01 and ***P < 0.001.

Immunoblotting analyses showed after 72 hours of SINEUP application in U-87 MG cells expressing *UBA5* pathogenic variants restored appropriate expression of UPR proteins, reducing p-PERK and p-eIF2α and increasing IRE1α (Fig. 7B). Target genes including BiP and GRP94 are also downregulated as a result, and PARP cleavage is also significantly reduced, similar to levels observed in UBA5^A371T/A371T^. Quantification of ER size also showed reduction in ER swelling in UBA5^A371T/R55H^ and UBA5^A371T/F292^ cells after SINEUP application (Fig. 7D). Together, these findings show that application of synthetic SINEUP constructs can modestly augment the expression of UBA5 and UBA5-UFM1 conjugates, thus rescuing the defects in ER homeostasis.

Next, we used CRISPRa-dCas9 method to increase *UBA5* gene expression as an alternative mechanism for rescue. We established UBA5^A371T/R55H^ and UBA5^A371T/F292*^ cells that stably express dCas9^-VPR^, where dCas9 is fused to a tripartite activator, VP64-p65-Rta (VPR) (*68*) (Fig. S9D), and designed 5 different sgRNA within the promoter element of *UBA5*. *UBA5* sgRNA were delivered via lentiviral transduction, and after 72 hours, all five sgRNA led to increased UBA5 expression when compared to the NTC construct in both compound heterozygous cell lines (Fig. 7E, 7F and Fig. S9E). At the protein level, we quantified a modest 2-fold increase for all sgRNA in both cell lines (Fig. 7F). It’s important to note that all sgRNA target *UBA5* genomic sequence, thus enhancing the expression of both isoforms, unlike SINEUP that is designed to target the processed mRNA of isoform 1. Moreover, we noted restoration of UPR protein expressions following augmented UBA5 expression, suggesting ER homeostasis is restored (Fig. 7E).

### Application of synthetic SINEUP increases abundance of UBA5 gene product and transiently rescues aberrant neuronal activity in patient-derived CO

Given that restoring *UBA5* expression in U-87 MG cells was able to rescue associated cellular defects, we then asked whether application of SINEUP in patient-derived CO can rescue aberrant neuronal firing. We delivered synthetic SINEUP constructs to 100-day old CO cultures vial lentiviral transduction and quantified a 1.5-fold increase in UBA5 (isoform 1) expression at 72 hours after transduction (Fig. 8A and 8B). We also observed rescue of UPR protein expression in proband CO following SINEUP application (Fig. 8A). Next, we measured the weighted mean firing rate of CO during a 2.5 week time span (Fig. 8C). On day 115, one CO was plated per well in an MEA plate and allowed to attach for the next 10 days. On day 125 (D-5), a recording was taken to establish baseline activity, then on day 130 (D0), SINEUP constructs were introduced, and another recording was taken immediately, then at D2, 3, 4 and 12 after SINEUP application. We observed a reduction in weighted mean firing rate for both proband CO between D2 to D4 after SINEUP application, in comparison to untreated CO or CO treated with NTC. Interestingly, we observed this rescue is only transient as SINEUP treated CO showed similar weighted mean firing rate as untreated or NTC at D12. Collectively, these findings suggest that increasing expression of the shared p.A371T allele may alleviate *UBA5*-associated disease phenotypes.

**Figure 8.**
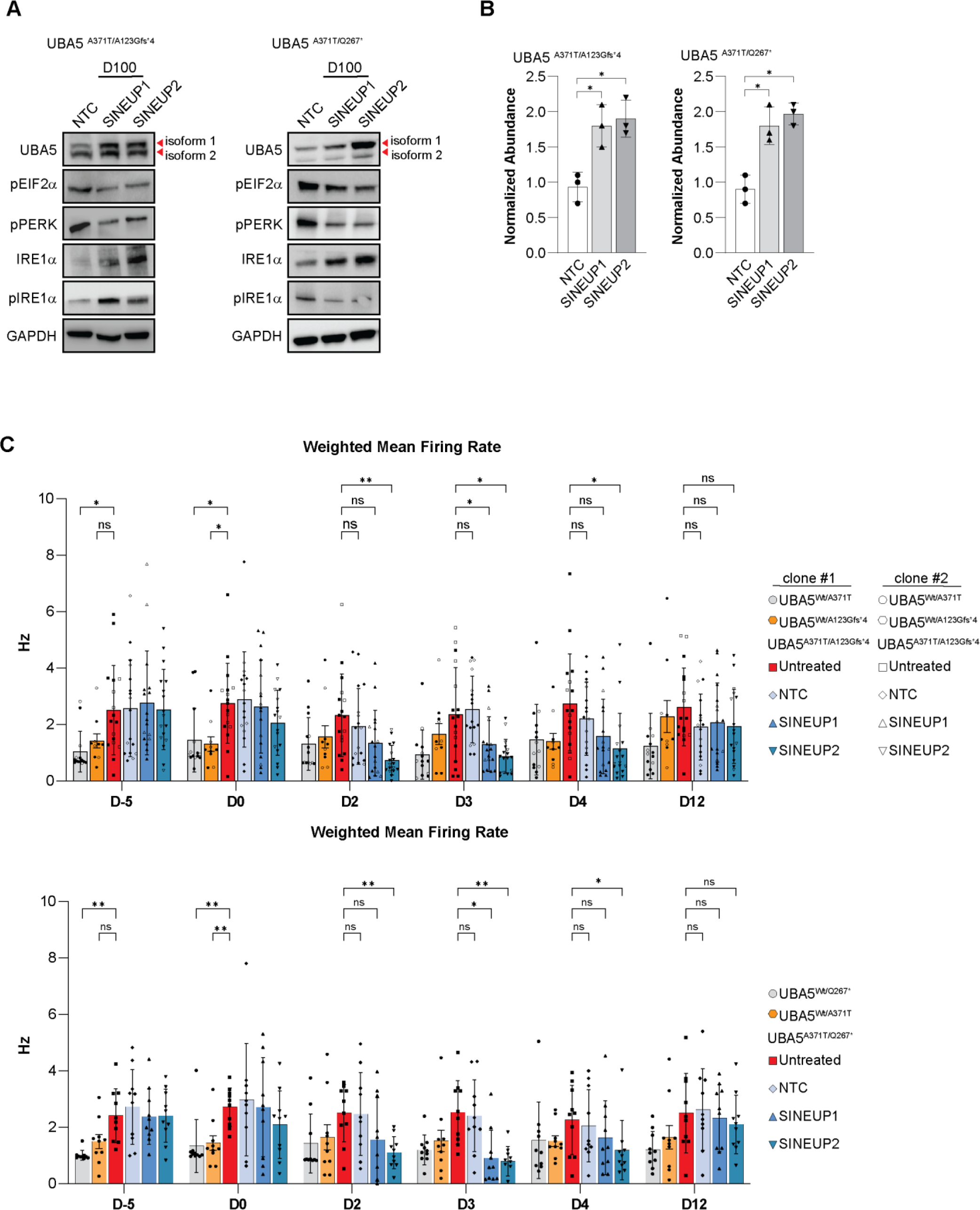
Genetic rescue of *UBA5* expression by SINEUP rescues electrophysiological defects in proband CO. **(A)** Immunoblot analysis restored expression UBA5 and UPR pathway in 100-day old CO derived from UBA5 probands after 72 hr of SINEUP treatment. NTC served as negative control for SINEUP. GAPDH served as loading control. All images shown in the manuscript are representative images of 3 independent experiments. **(B)** Quantification of UBA5 protein abundance following SINEUP treatment, first normalized to GAPDH, and then to averaged NTC treatment. Each data point represents one experiment, plotted as mean ± SD. *P<0.05. **(C)** Weighted mean firing rate of proband CO as measured by MEA. The same set of 125-day old CO were measured 5 days prior (D-5) to SINEUP treatment (D0), then measured at D2, D3, D4 and D12 after SINEUP treatment. Each data point represents one CO, plotted as mean ± SD. *P<0.05, **P<0.01 and ns: not significant.

## Discussion

The clinical phenotypes presented in *UBA5*-associated DEE most strikingly affect the brain (*1–8*), despite *UBA5* being ubiquitously expressed in all tissues and during all stages of life. Germline knockout of *Uba5* in mice results in embryonic lethality (*27*) and there are no known individuals with UBA5-associated DEE with biallelic null variants. A recent study utilized humanized *Drosophila* models to assess the molecular strength of patient variants and was able to empirically classify the severity of disease-causing alleles, but did not investigate neuronal defects (*40*). Thus, there is still no patient-derived disease model to investigate development of the brain, assess clinical phenotypes and test potential therapy. In this study, we identified two patients with previously unpublished nonsense *UBA5* variants on one allele (p.Q267* and p.A123Gfs*4) and a recurrent hypomorphic variant on the other allele (p.A371T) that presented with *UBA5*-associated DEE phenotypes. We established cortical organoid models of these patients, along with healthy parental controls to investigate the human-specific pathogenesis of UBA5^A371T/Q267*^ and UBA5^A371T/A123Gfs*4^. We applied unbiased bulk and single-cell RNAseq analyses that identify significant reduction in GABAergic inhibitory processes. Our scRNAseq analysis uncovered a striking deficit in both progenitors and mature inhibitory GABAergic interneurons.

We noted a significant reduction of GABAergic markers *GAD1* and *GAD2*, which encode the enzymes that convert glutamate to GABA; expression of *GAD2* is necessary for promoting the differentiation of GABAergic interneurons (*69, 70*). This suggests that the loss of GABAergic neuron population observed in CO derived from *UBA5* proband may be due to insufficient expression of *GAD1* and *GAD2* during differentiation. Interestingly, we noted a more significant reduction in the expression of *GAD2* compared to *GAD1* in proband CO (Fig. 2E). In mature GABAergic interneurons, GAD1 and GAD2 perform distinct functions within the same cell. GAD1 synthesizes GABA for tonic, non-vesicle neurotransmitter release, whereas GAD2 is localized in the axon terminals on membranes and synaptic vesicles, for activity-dependent GABA synthesis and vesicle release of GABA during intense synaptic activities (*71, 72*). In this regard, decreased *GAD2* expression may further contribute to impairment in GABA synaptic release and inhibitory GABA function in the limited number of mature GABAergic interneurons present in *UBA5* proband CO. Thus, reduced *GAD2* expression can first affect interneuron differentiation and then affect synaptic GABA release in the remaining mature interneurons. These findings all show changes in the excitatory-inhibitory balance (*73, 74*). In support of this, we observed augmented excitability in proband CO, with increased firing rate but decreased network activity. GABAergic interneurons counteract excitation through local circuitry that connects excitatory and inhibitory neurons, and impaired interneuron function can lead to epilepsy (*75, 76*). Loss of GABAergic interneurons can also account for the smaller proband CO size. Interestingly, progressive microcephaly is also a clinical feature DEE caused by biallelic mutations of *UFC1* or *UFM1* (*28*). More detailed analysis of corticogenesis is vital to understand how *UBA5* pathogenic variants lead to microcephaly and epilepsy in patients. Moreover, pinpointing the onset and evolution of aberrant neurodevelopment will provide information about the most effective therapeutic window(s). The most effective approach to treatment may require fetal therapy, similar to *in utero* enzyme-replacement therapy to successfully treat infantile-onset Pompe’s disease (*77*).

Our protocol generated CO that contained GABAergic interneurons that originated from the caudal ganglion eminence (CGE), evident by high expression of *CALB*, and low expression of *PV* and *SST* (Fig. 2E and Fig. S4). This group accounts for 30-40% of GABAergic cortical neuron population, whereas the rest are originated from the medial ganglion eminence (MGE) (*78*). Interneurons from both CGE and MGE differ in their axonal targets, morphology and firing patterns (*78*). Thus, to better understand how *UBA5* pathogenic variants affect corticogenesis, it would be necessary to investigate differentiation of GABAergic interneuron in the MGE using other organoid models.

At the molecular level, we confirmed U-87 MG cells expressing *UBA5* pathogenic variants (UBA5^A371T/R55H^ and UBA5^A371T/F292*^) exhibit impaired ufmylation pathway function shown by decreased UBA5-UFM1 conjugates. The UBA5-UFM1 conjugation was shown to be the rate-limiting step in the ufmylation pathway that determines pathway activity (*79*). We then showed *UBA5* pathogenic variants perturbed ER homeostasis, leading to sustained activation of the UPR pathway, including PERK and ATF6. Regulation of protein synthesis, folding and quality control is important for synaptic plasticity in the adult brain. During long-term potentiation and depression, acute changes in protein synthesis are necessary to modulate the efficacy of synaptic connections (*80, 81*). In the developing brain, protein synthesis is much higher in progenitors to meet the demands of proliferating and maturing neurons, and UPR pathway is involved in different aspects of neurogenesis (*82–86*). For example, exacerbated PERK-eIF2α signaling interferes with the generation of intermediate progenitors and cortical layering, leading to microcephaly (*85*). ATF6-directed transcription also contributes to neurodevelopment (*82*). Thus, in *UBA5*-associated DEE, exacerbated UPR pathway activity may result in insufficient protein production, such as GAD1 and GAD2 during neuronal development that leads to loss of interneurons. Post development, neuronal activity dependent GAD2 protein production may also be hindered, preventing proper interneuron response. Collectively, these findings all highlight the importance for both temporal and spatial regulation of the UPR pathway. While there’s no doubt that pathogenic variants of *UBA5* lead to neurodevelopmental disorders (*1–8, 28*), there are still important outstanding questions that need to be addressed to better understand disease manifestation and highlight potential therapeutic avenues. How is the expression of UBA5 and activity of the ufmylation pathway regulated during neurogenesis? Are there additional perturbations to gene expression besides *GAD1* and *GAD2* that affect neurodevelopment in *UBA5*-associated DEE? We observed exacerbated UPR pathway activity in cells expressing *UBA5* pathogenic variants, but how does impaired ufmylation pathway lead to increased PERK and ATF6 activation, and what are the substrates involved? Can we target so-said substrates of the ufmylation pathway or UPR pathway as potential therapeutic outlets?

The observation of healthy individuals with homozygous p.A371T allele is compelling, offering a unique therapeutic avenue for the >75% of *UBA5* patients that all share this common hypomorphic allele (*5, 7, 8*). Furthermore, if not p.A371T, it is likely that all affected individuals have at least one hypomorphic allele, so upregulation of transcription or translation could be effective for all affected individuals (*4, 6–8, 40*). We demonstrated that cells expressing two copies of the p.A371T allele function nearly identically to those expressing wildtype *UBA5*, with no alterations to UPR pathway activities. We utilized two modalities to upregulate the expression of the p.A371T variant, first using synthetic SINEUP to increase translation and then using CRISPRa-dCas9^VPR^ to increase transcription. SINEUP offered the more modest level of upregulation compared to CRISPRa-dCas9^VPR^, which is therapeutically important for *UBA5*-associated DEE and other similar haploinsufficient disorders (*87*). Overexpression of UBA5 is detrimental to proper ufmylation, mimicking *UBA5* deletion with loss of UBA5-UFM1 conjugation (*88*). This could explain why we only observed a transient rescue of neuronal firing in proband CO treated with SINEUP. As a proof of concept, we packaged SINEUP in a lentiviral plasmid for ease of delivery, which could result in overexpression of UBA5 following genome integration of SINEUP. Indeed, we observed overexpression of UBA5 at 9 days following SINEUP application, with >3.5 fold increase. It would be beneficial to explore delivery of SINEUP via other delivery methods including adeno-associated virus and lipid nanoparticles (*89–92*). These delivery vehicles do not allow for genome integration and have been approved for human trials by the FDA (*89–92*).

It is important to note that there are other modalities regulating gene expression that can be utilized to boost UBA5 abundance, and it would be beneficial to assess their utility. Tethered mRNA Amplifier utilizes the 3’UTR and enhances transcript stability to promote translation (*93*). Similarly, translation-activating RNA is a bifunctional RNA-based molecule that brings together target mRNA and translation machinery to enhance protein output (*94*). All of these technologies, including SINEUP and CRISPRa, are mutation agnostic and can be broadly considered for any haploinsufficient disorders. However, detailed investigation of delivery methods, efficacy and outcome is warranted. Collectively, our findings outline the first patient-derived neuronal model for investigating *UBA5*-associated DEE and we were able to identify defects in corticogenesis with regards to GABAergic interneuron differentiation. *UBA5* pathogenic variants perturb ER homeostasis by exacerbating UPR pathway activity. We characterized the hypomorphic p.A371T allele and showed cells expressing two copies of p.A371T functions similarly to the wildtype. We took advantage of this finding and boosted the expression of the one copy of p.A371T allele in patient-derived cells using SINEUP and CRISPRa-dCas9^VPR^, rescuing various disease-associated defects. Our study illustrates the importance of disease modeling using patient derived samples and highlighted a potential therapeutic intervention that can be mutation agnostic.

## Materials and methods

### Human subjects

All protocols and consenting procedures were approved by the IRBs of University of Washington and St. Jude Children’s Research Hospital. Informed consent was obtained from the parents or legal guardians of all patients and adult parental control participants consented for themselves. Skin biopsies from UBA5 patients and healthy parental controls were collected according to standard collection procedures by a qualified healthcare professional.

### Generation and maintenance of cortical organoids (CO)

Cortical organoids (CO) were generated by modifying a dorsal forebrain differentiation protocol (*95*). iPSC lines were dissociated to single cell using Accutase (STEMCELL Technologies). Cells were then aggregated into embryoid bodies (EB) at a density of 10,000 cells/well in 96-well v-bottom low attachment plates. DMEM-F12 (Gibco) based EB medium included 20% Knock-out Serum (Gibco), 3% ES-FBS (Gibco), 0.1 mM β-Mercaptoethanol (Gibco), 5 µM SB-431542 (Tocris), 2 µM Dorsomorphin (Sigma Aldrich) and 3 µM IWR1-endo (Millipore). Media was supplemented with 20 µM Y-27632 for the first week. From day 4 to day 16, CO were cultured in GMEM-based neural induction medium with 20% Knock-out Serum, 5 µM SB-431542 and 3 µM IWR1-endo. From day 18 to day 28, media was supplemented with 1 N2 (Gibco), 1X B-27 without Vitamin A (Gibco), 1X CD Lipid Concentrate (Gibco), 20 ng/mL human-recombinant FGF2 (PeproTech) and 20 ng/mL human-recombinant EGF (PeproTech). On day 22, CO were transferred to bioreactors spinning at 40 rpm. From day 30 to day 50, media was supplemented with 10% ES-FBS, 1X N2, 1X B-27 without Vitamin A, 1X CD Lipid Concentrate and 5 µg/mL Heparin (Sigma Aldrich). From day 50 and onwards, CO were maintained in BrainPhys Neuronal Media with 10% ES-FBS, 1X N2, 1X B-27 without Vitamin A, 10 ng/mL human-recombinant brain-derived neurotrophic factor, BDNF (PeproTech) and 10ng/mL human-recombinant glial cell-derived neurotrophic factor, GDNF (PeproTech).

### Generation of U-87 MG cells with UBA5 pathogenic variants

U-87 MG *UBA5* genetically modified cell lines were generated using CRISPR technology in the Center for Advanced Genome Engineering (St. Jude). All cell lines were maintained at 37°C with 5% CO2 and passaged using 0.05% Trypsin (Gibco) when reaching 70–90% confluency. Cells were grown in DMEM supplemented with 10% fetal bovine serum (Gibco), 2 mM L-glutatmax (Gibco), 100 U/mL antibiotic-antimycotic (Gibco). Additional details are available in Supplemental Methods, Supplemental Table 2.

### Cortical organoid single cell capture and scRNAseq

CO were collected for single cell capture (N=3) at 100 days after initial seeding of embryoid body. CO were dissociated using papain solution containing DNase (Worthington). CO were washed 3 times using PBS, and then cut up using Vannas Spring Scissors (Fine Science Tools). Samples were then incubated in 2 mL papain solution at 37°C for 1 hr, and samples were triturated by manually pipetting every 20 mins during the 1 hr incubation period. Dissociated cells were then filtered through a 40 µm cell strainer, and centrifuged at 300G for 5 mins, papain removed and resuspended in 1% FBS in PBS. Using the Chromium Single Cell 3’ Reagent Kits v3.1 (10X Genomics), 4000-7000 cells were captured per lane, single-cell libraries were sequenced on a NovaSeq S2 flow cell.

### 10X Genomics CellRanger software and data filtering

Illumina 10X Genomics scRNAseq base-call files were converted to FASTQ, which aligned to the hg38 genome and transcriptome using the CellRanger v3.0.2 pipeline, for a gene–cell expression matrix. The count matrices of 4 samples (2 proband CO and 2 control CO) were processed for filtering and downstream analysis using Partek Flow software. In short, the quality controls for cell filtering were performed on the gene expression matrices as follows: 1) cells with fewer than 1,000 or more than 20,000 unique molecular identifiers were removed; 2) cells expressing less than 500 or larger than 5,000 unique genes were considered outliers and discarded; and 3) cells with a mitochondrial transcript proportion higher than 20% were filtered out. In addition, the genes with maximum count of 1 among all cells were excluded for further analysis. We used a commonly applied normalization approach count-per-million (CPM) normalization followed by log1p transformation. All features were subjected to principal component analysis (PCA), calculating top 100 principal components (PCs) by variance. The top 30 PCs were used for graph-based clustering using Smart Local Moving (SLM) algorithm and visualized in Uniform Manifold Approximation and Projection (UMAP). Finally, the biomarkers were identified by computing the features that were expressed highly when comparing each cluster.

### Bulk RNA sequencing of CO

CO (n=3) were pooled together for RNA extraction using *Quick*-RNA Miniprep Kit (Zymo Research) according to the manufacturer’s instructions. Libraries were prepared from total RNA with the TruSeq Stranded mRNA Library Prep Kit according to the manufacturer’s instructions (Illumina). Paired-end 100-cycle sequencing was performed on a NovaSeq 6000 (Illumina). The 100-bp paired-end reads were trimmed, filtered against quality (Phred-like Q20 or greater) and length (50-bp or longer), and aligned to a human reference sequence GRCh38/hg38 by using CLC Genomics Workbench v20 (Qiagen). The TPM (transcript per million) counts were generated from the CLC RNA-Seq Analysis tool. The differential gene expression analysis was performed by using the non-parametric ANOVA using the Kruskal-Wallis and Dunn’s tests on log-transformed TPM between biological replicates from proband CO and control CO, implemented in Partek Genomics Suite v7.0 software (Partek Inc.). The cutoff for significance is p < 0.01 and |log2R| > 1 (two-fold change) between two experimental sets. The gene sets enrichment and pathway analysis were performed using GSEA (https://www.gsea-msigdb.org/).

### Multielectrode array reading and analysis

CO were plated per well in Matrigel-coated 24-well CytoView plates (Axion Biosystems), media was half-changed every 4 days with 2 µg/mL Natural Mouse Laminin (ThermoFisher Scientifics). Recordings were performed starting on day 115 using a Maestro Edge MEA system and AxIS Software Spontaneous Neural Configuration and Viability Module. Cytoview plates were maintained with 5% CO2 at 37°C during recording. CO were equilibrated for 5 mins in the MEA system, and then >10.5 mins of data were recorded. Spikes were detected with AxIS software using an adaptive threshold crossing set to 5.5 times the standard deviation of the estimated noise for each electrode (channel). For the MEA analysis, the electrodes that detected at least 5 spikes/min were classified as active electrodes using Axion Biosystems’ Neural Metrics Tool. Bursts were identified in the data recorded from each individual electrode using an inter-spike interval (ISI) threshold requiring a minimum number of five spikes with a maximum ISI of 100 ms. At least ten spikes under the same ISI with a minimum of 25% active electrodes were required for network bursts in the well.

### Statistics

All graphs display the mean ± SD and each data point representing a unique sample or experiment as indicated. Statistical analysis was performed by 2-way ANOVA with Tukey’s multiple comparisons test, with P value indicated as *P < 0.05, **P < 0.01, ***P < 0.001, ****P < 0.0001 and ns: not significant.

## List of Supplemental Materials

### Materials and Methods

Fig S1-S9

Table S1-S5

References (96–98)

## Author Contributions

HC and HCM conceived and designed the study. YDW performed data analysis for bulk and single-cell RNAseq. HC acquired data, performed experiments, and analyzed all data, with help from AWB. EAF and ESB organized patient data and consent. RB and SPM designed and generated CRISPR-edited U-87 MG cells. HC and HCM wrote the manuscript with input from all authors.

## Acknowledgements

We thank the patients and their families for donating materials to make these studies possible. We thank members of the Mefford lab for helpful discussions and critical reading of the manuscript, and Dr. Alex Carisey (SJCRH), James Messing (SJCRH), Dr. Aaron Taylor (SJCRH), Dr. Aaron Pitre (SJCRH) and Dr. Nicolas Denan (SJCRH) for their valuable time and input into experimental design. We thank Dr. Danny Miller (U. Washington) for performing targeted long-read sequencing and Jennifer Dempsey (U. Washington) for assistance with clinical records. We thank Dr. Anjana Nityanandam (SJCRH) and Kyle Newman (SJCRH) for helping to develop CO protocol and consulting CO experiments. We would like to also acknowledge the St. Jude Children’s Research Hospital Vector Development & Production Core, Hartwell Center, Center for Advanced Genome Engineering, Center for Modelling Pediatric Diseases and Cell and Tissue Imaging Center-Light Microscopy (supported by NCI P30 CA021765).

## Competing interests

The authors have declared that no conflict of interest exists.

## Data and materials availability

All data associated with this study are present in the paper or the Supplementary Materials. Sequencing data have been uploaded to GEO.

## Notes

### Competing Interest Statement

The authors have declared no competing interest.

