## Supplemental Figures for "Patient derived model of *UBA5-*associated encephalopathy identifies defects in neurodevelopment and highlights potential therapies"

### Supplemental Figure 1

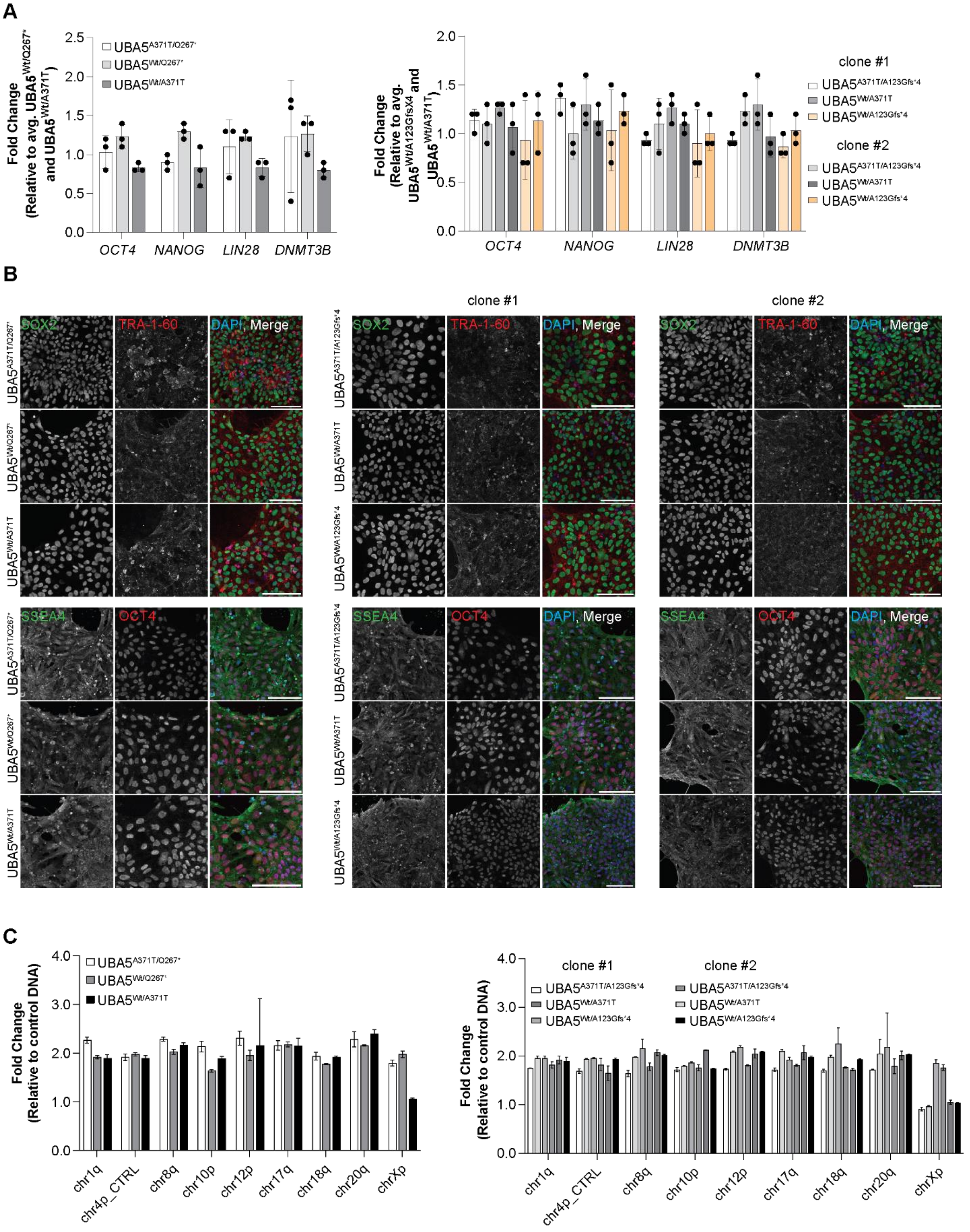

**Supplemental Figure 1. Characterization of iPSC reprogrammed from *UBA5* probands and healthy parental controls. (A)** Transcript level of key pluripotency markers in all iPSC lines. Each data point represents one experiment, plotted as mean ± SD. **(B)** Representative images of immunofluorescent staining of key pluripotency markers. Scale bars: 100 µm. **(C)** Analysis of eight most common karyotypic abnormalities as detected by the hPSC Genetic Analysis Kit™ (StemCell Technologies™), plotted as mean ± SD.

### Supplemental Figure 2

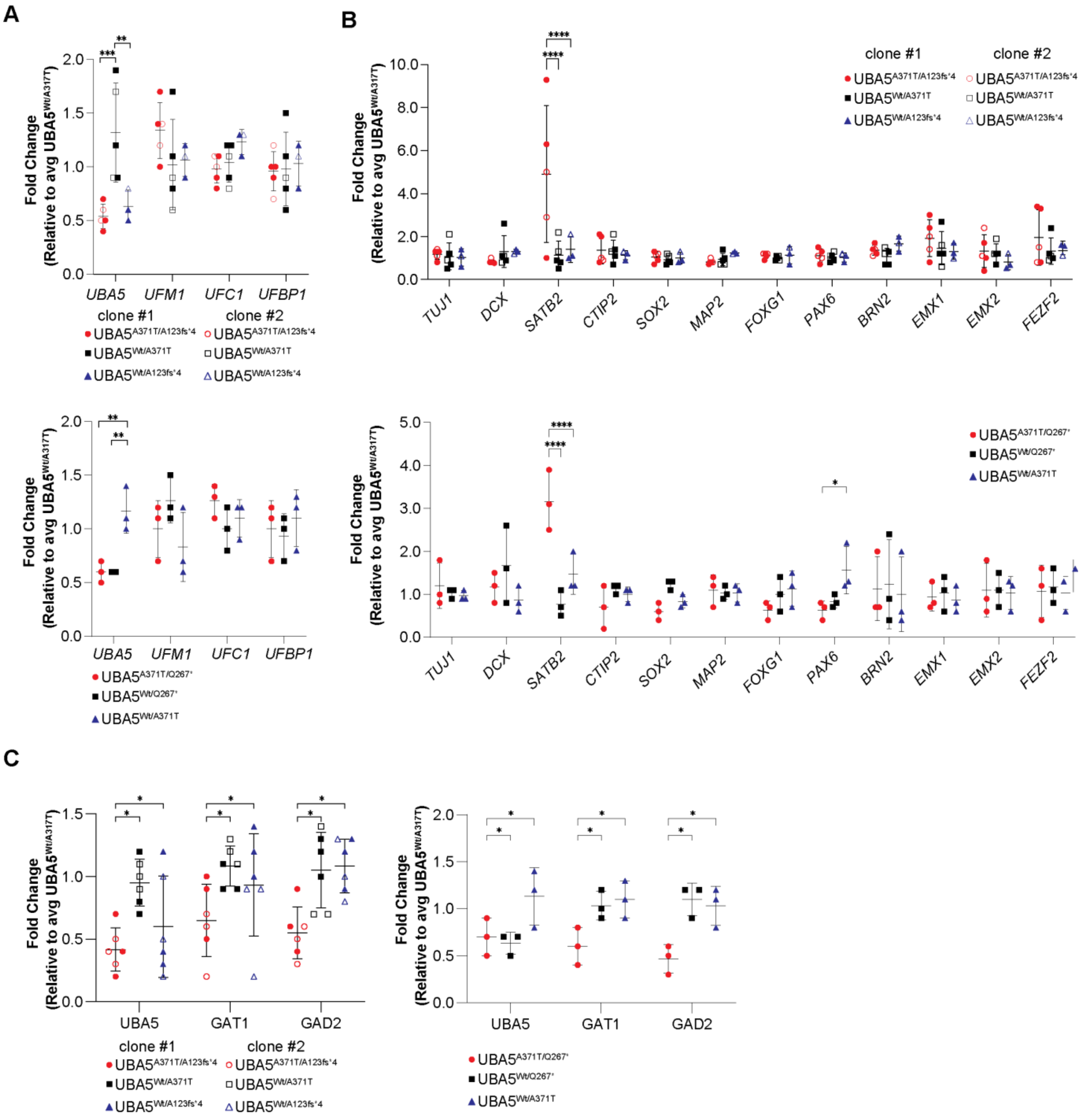

**Supplemental Figure 2. Expression of ufmylation and neuronal markers in 100-day old CO. (A)** Transcript level of ufmylation pathway proteins, normalized to averaged UBA5$^{Wt/A371T}$. Each data point represents three CO, plotted as mean ± SD. **P < 0.01 and ***P < 0.001. **(B)** Transcript level of various neuronal cell type markers, normalized to averaged UBA5$^{Wt/A371T}$. Each data point represents three CO, plotted as mean ± SD. *P < 0.05 and ****P < 0.0001. **(C)** Quantification of protein abundance, first normalized to GAPDH, and then to averaged UBA5$^{Wt/A371T}$. Each data point represents one experiment, plotted as mean ± SD. *P<0.05.

### Supplemental Figure 3

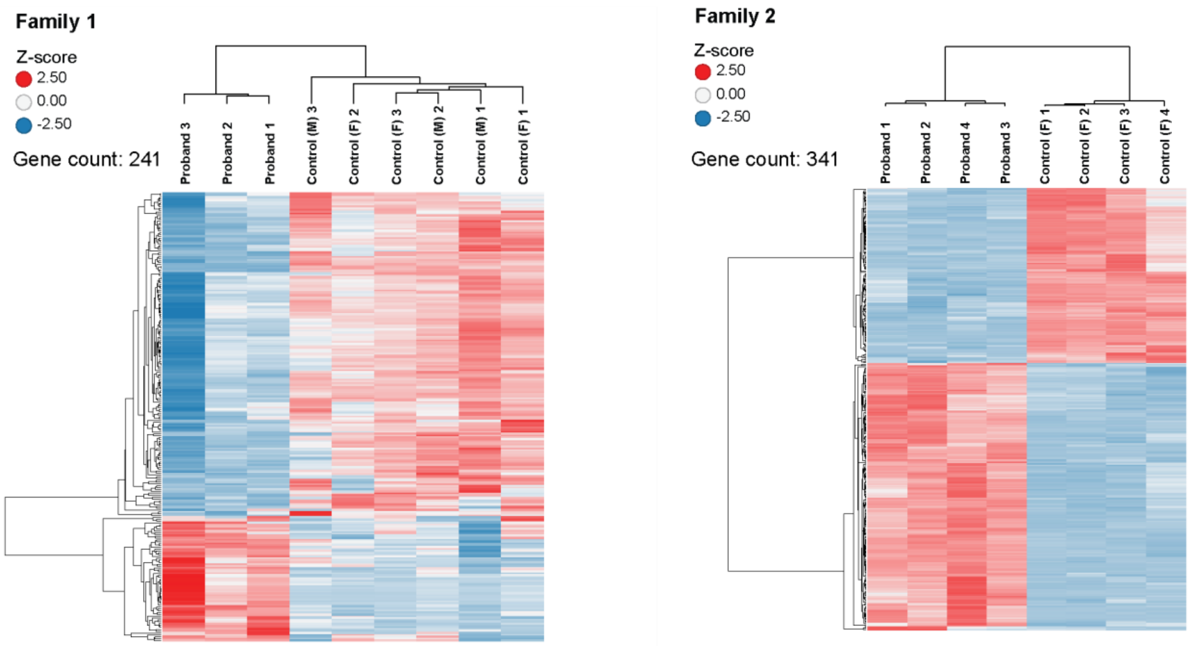


**Supplemental Figure 3. Heatmap of differentially expressed genes in bulk RNAseq analysis of 100-day old CO.** Heatmap of expression of the top differentially expressed genes based on hierarchical clustering for families 1 (UBA5$^{A371T/A123Gfs*4}$) and 2 (UBA5$^{A371T/Q267*}$). Control M: paternal control; Control F: maternal control.

### Supplemental Figure 4

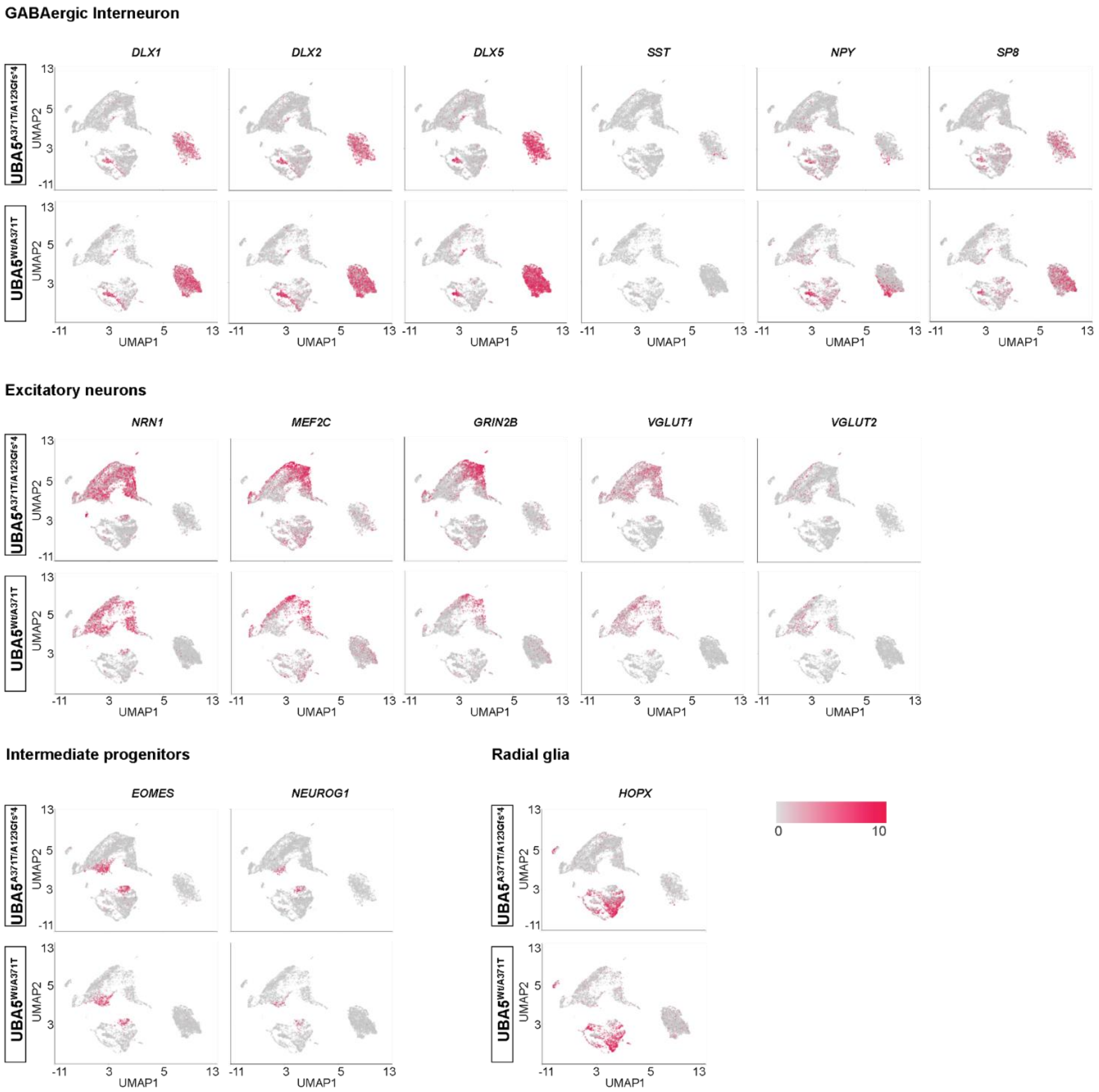


**Supplemental Figure 4. Feature plots showing gene expression differences between *UBA5* proband and control CO from scRNAseq analysis.** Comparison of specific marker expression between *UBA5* proband and control CO from scRNAseq analysis.

### Supplemental Figure 5

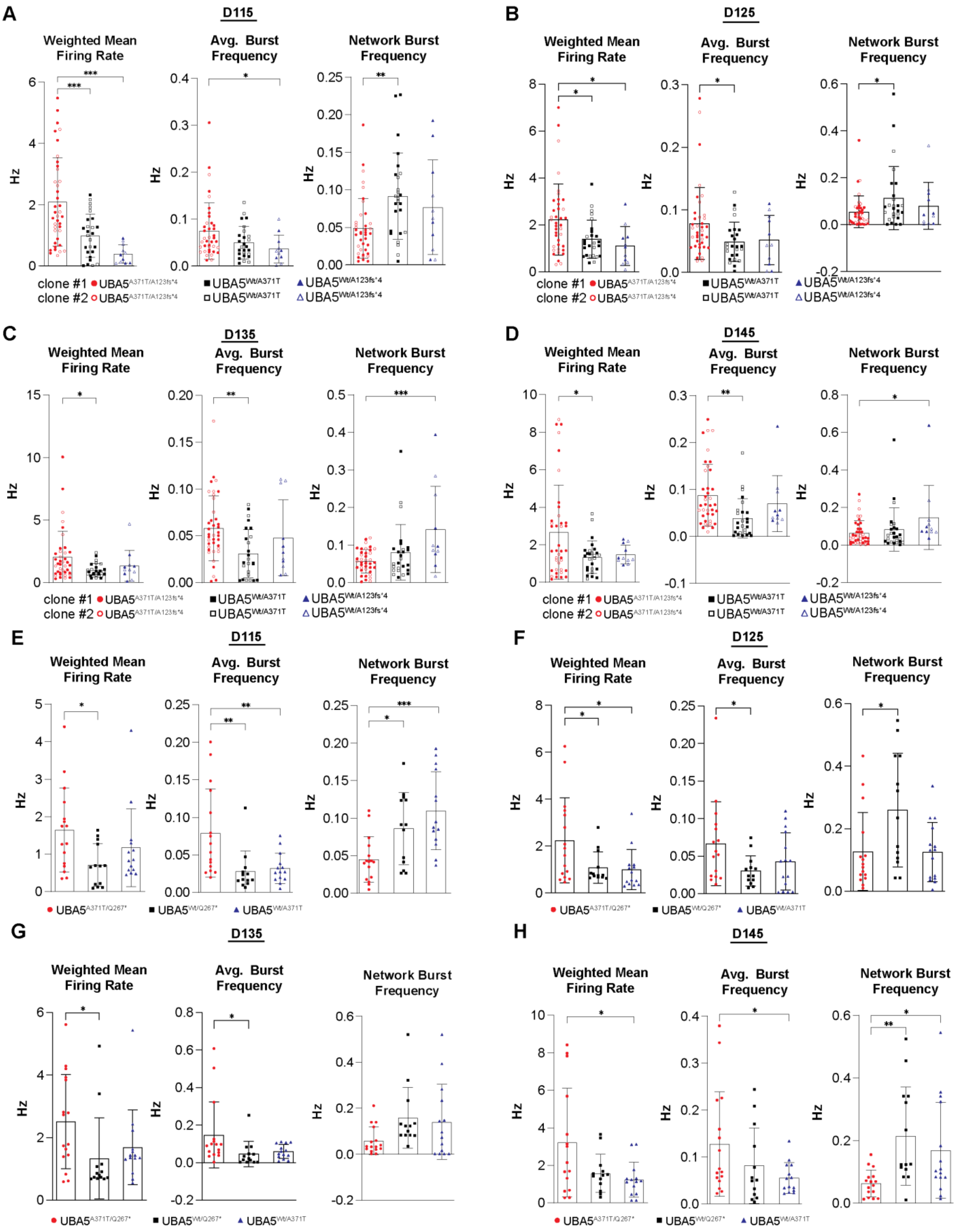

**Supplemental Figure 5. CO derived from *UBA5* probands showed aberrant network activity.** Functional characterization of CO derived from *UBA5* probands and controls using MEA at **(A and E)** D115, **(B and F)** D125, **(C and G)** D135 and **(D and H)** D145, showing changes in weighted mean firing rate, averaged burst frequency and network burst frequency compared to controls. Each data point represents one CO, plotted as mean ± SD. *P < 0.05, **P < 0.01 and ***P<0.001.

#### Supplemental Figure 6

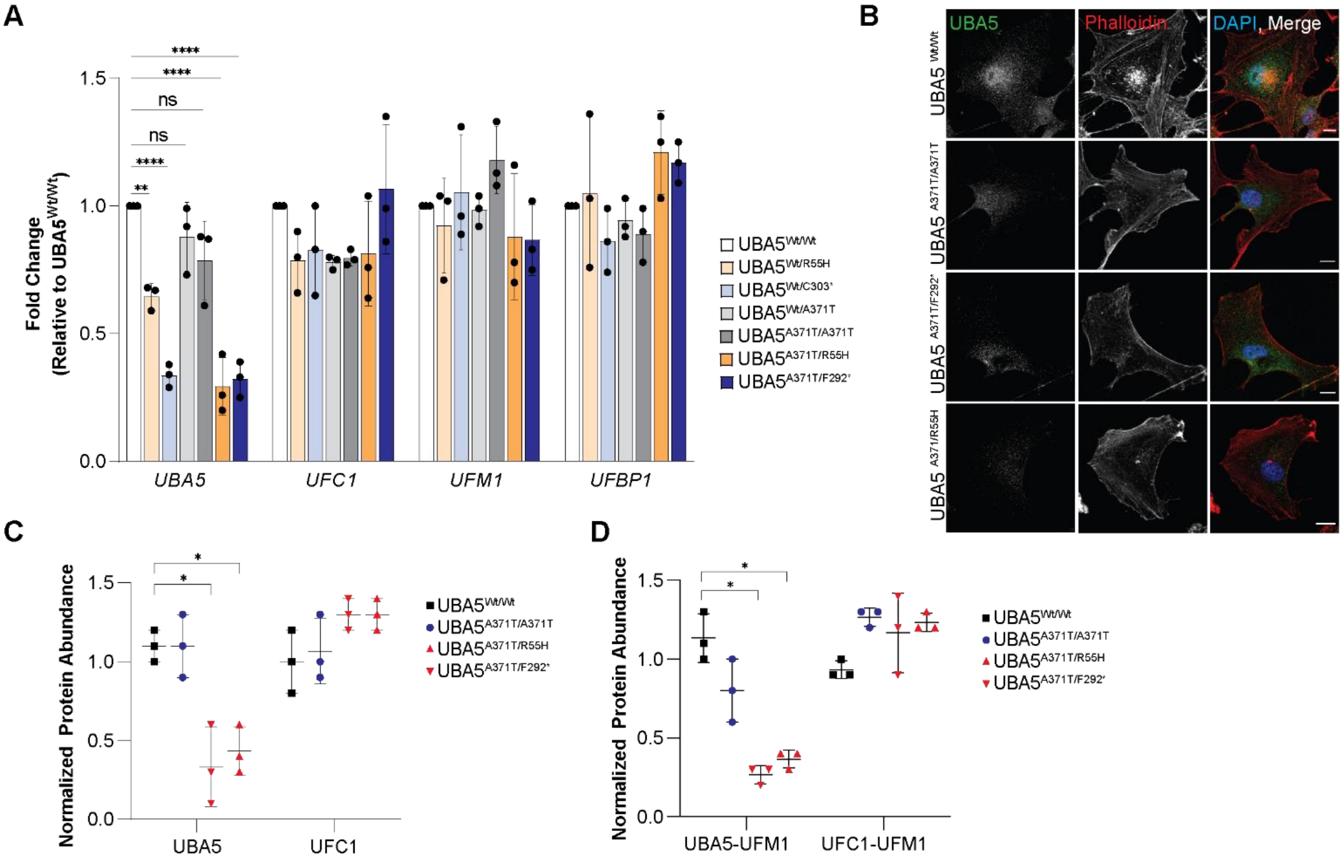


**Supplemental Figure 6. Characterization of U-87 MG cells expressing *UBA5* pathogenic variants. (A)** Transcript level of ufmylation pathway proteins, normalized to averaged UBA5$^{Wt/Wt}$. Each data point represents one experiment, plotted as mean ± SD. **P < 0.01 and ****P < 0.0001. **(B)** Representative images of immunofluorescent staining of UBA5 and phalloidin. Scale bars: 10 µm. **(C)** Quantification of protein abundance, first normalized to GAPDH, and then to averaged UBA5$^{Wt/Wt}$. Each data point represents one experiment, plotted as mean ± SD. *P<0.05. **(D)** Quantification of protein abundance, first normalized to GAPDH, and then to averaged UBA5$^{Wt/Wt}$. Calculated values include samples with and without DTT. Each data point represents one experiment, plotted as mean ± SD. *P<0.05.

### Supplemental Figure 7

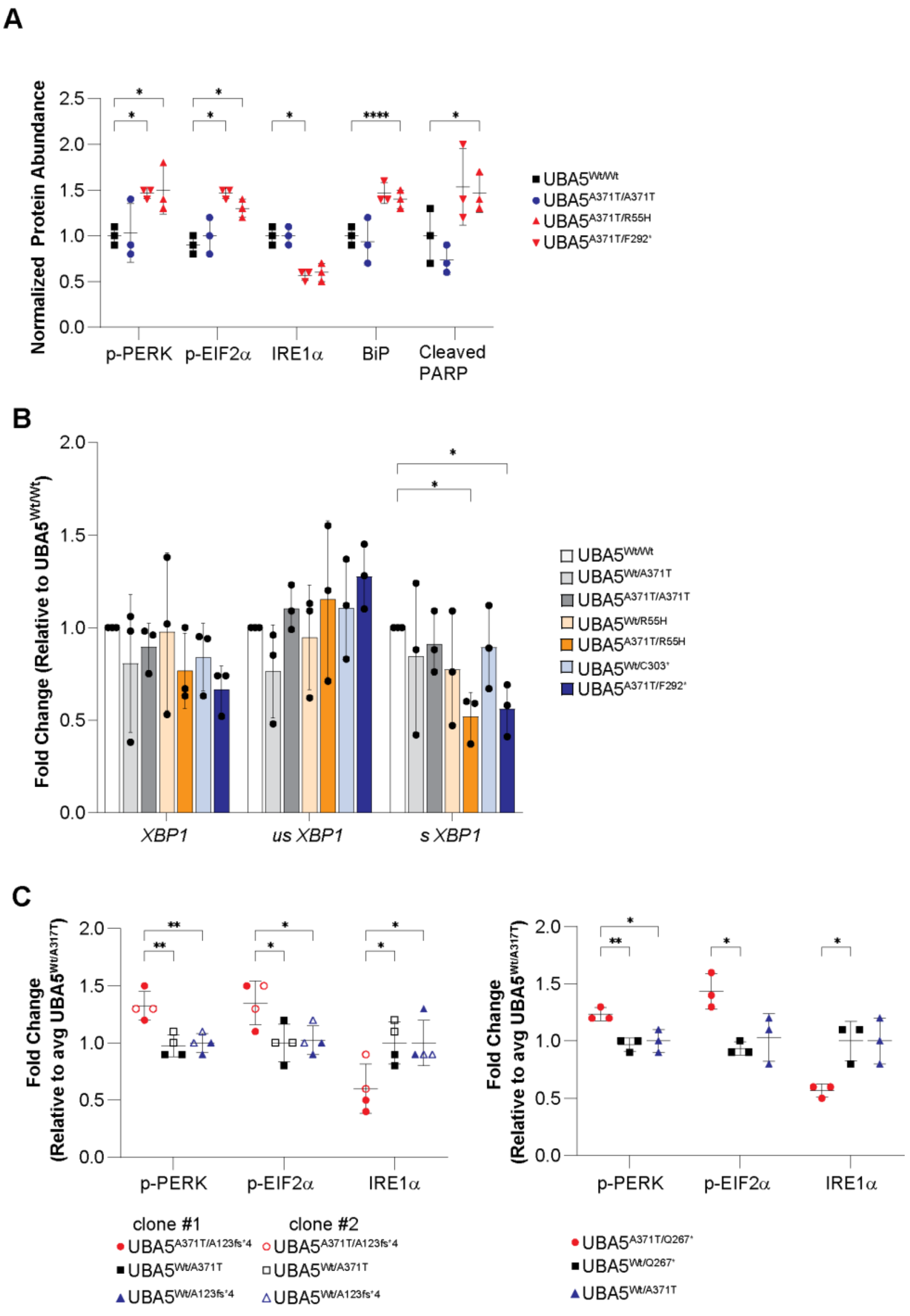


**Supplemental Figure 7.** ***UBA5* pathogenic variants disturb ER homeostasis in U-87 MG cells and CO. (A)** Quantification of protein abundance, first normalized to GAPDH, and then to averaged $UBA5^{Wt/Wt}$. Each data point represents one experiment, plotted as mean ± SD. *P<0.05 and ****P<0.0001.**(B)** Transcript level of total *XBP1*, spliced (s) and unspliced (us) *XBP*, normalized to $UBA5^{Wt/Wt}$. Each data point represents one experiment, plotted as mean ± SD. *P < 0.05. **(C)** Quantification of protein abundance, first normalized to GAPDH, and then to averaged $UBA5^{Wt/A371T}$. Each data point represents one experiment, plotted as mean ± SD. *P<0.05 and **P<0.01.

### Supplemental Figure 8

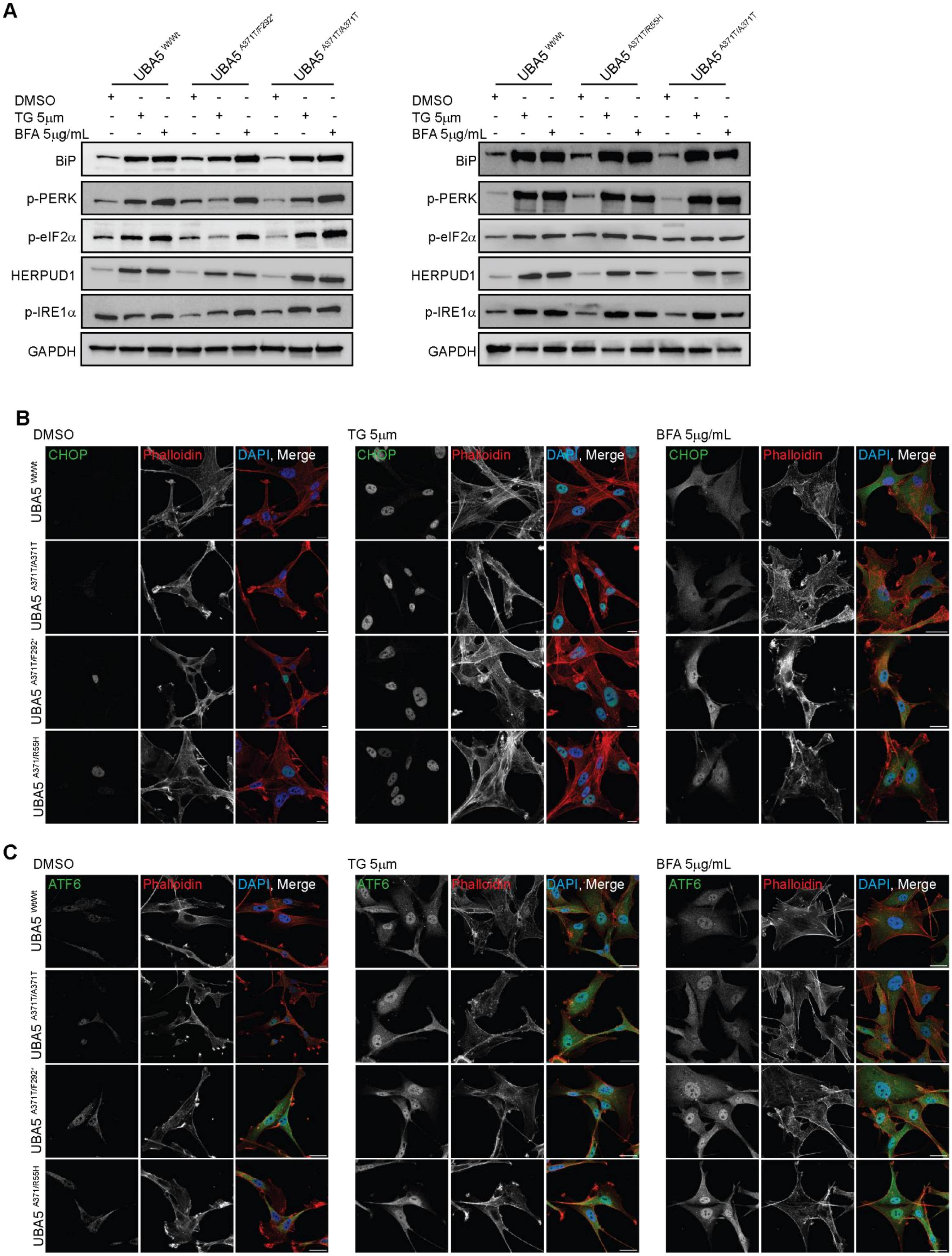

**Supplemental Figure 8.** ***UBA5*** **pathogenic variants in U-87 MG cells do not alter ER stress response induced by small molecules. (A)** Immunoblot analysis showed an increase of various components of the UPR response pathways in U-87 MG cells with pathogenic *UBA5* variants following 6 hr of TG or BFA treatment. DMSO served as vehicle control. GAPDH served as loading control. **(B-C)** Representative images of immunofluorescent staining with **(B)** CHOP and phalloidin, **(C)** ATF6 and phalloidin in U-87 MG cells with *UBA5* pathogenic variants following 6 hr of TG or BFA treatment, demonstrating nuclear localization of ATF6 and CHOP in all cell lines. DMSO served as vehicle control. Scale bars: 10 μm.

### Supplemental Figure 9

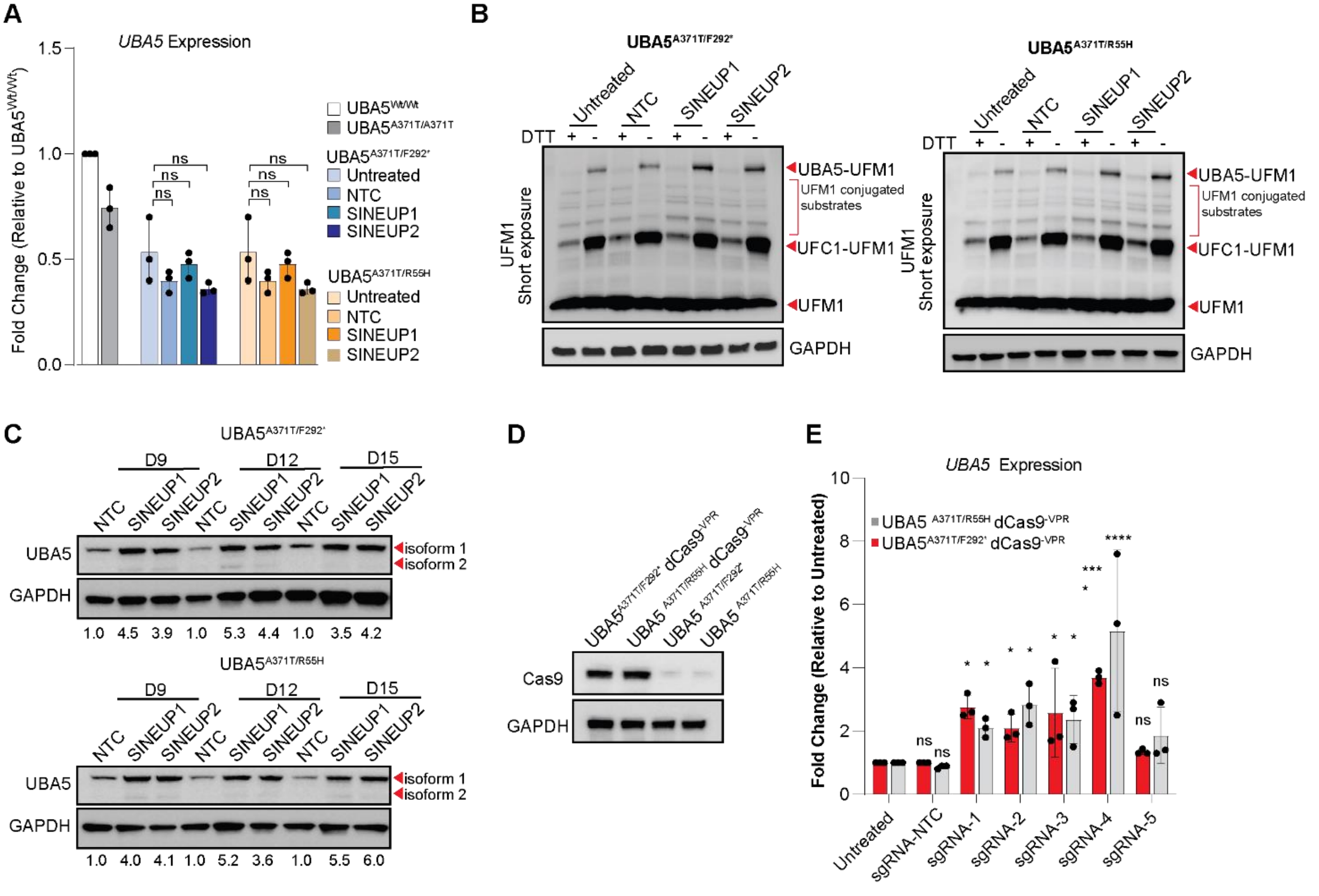


**Supplemental Figure 9. Quantitation of** ***UBA5*** **expression following SINEUP or CRISPRa treatment in U-87 MG cells. (A)** *UBA5* transcript levels remain unchanged in U-87 MG cells with *UBA5* pathogenic variants following 72 hr of SINEUP treatment. NTC served as negative control for SINEUP. Each data point represents one experiment, plotted as mean ± SD. ns: not significant. **(B)** Immunoblot analysis of UFM1 conjugates following 72 hr of SINEUP treatment, with and without reducing agent, DTT. NTC served as negative control for SINEUP. GAPDH served as loading control. All images shown in the manuscript are representative images of 3 independent experiments. **(C)** Immunoblot analysis showing augmented UBA5 expression 9 and 15 days after SINEUP treatment. NTC served as negative control for

SINEUP. Average UBA5 fold-change from three different experiments are as indicated. UBA5 level is first normalized to GAPDH, and then to averaged NTC treatment. GAPDH served as loading control. All images shown in the manuscript are representative images of 3 independent experiments. **(D)** Immunoblot analysis confirming stable expression of Cas9 in UBA5$^{A371T/F292*}$ dCas9$^{-VPR}$ and UBA5$^{A371T/R55H}$ dCas9$^{-VPR}$ cells. GAPDH served as loading control. All images shown in the manuscript are representative images of 3 independent experiments. **(E)** *UBA5* transcript levels increased in U-87 MG cells with pathogenic *UBA5* variants expressing dCas9-VPR following 72 hours of sgRNA treatment, compared to untreated. Each data point represents one experiment, plotted as mean ± SD. *P < 0.05, ***P<0.001 and ns: not significant.

**Supplemental Methods**

**Generation, characterization, and maintenance of iPSC lines**

Fibroblasts from *UBA5* patients and healthy parental controls were reprogrammed using the CytoTune-iPS 2.0 Kit (Thermo Fisher Scientifics) following manufacturer's instructions. iPSC were clonally isolated and maintained on feeder for 10 passages, then transferred to feeder-free Matrigel (BD Biosciences) coated plates and maintained in mTeSR1 (Stem Cell Technologies) and passaged at 70% confluence using ReLeSR (STEMCELL Technologies) with Y-27632 Dihydrochloride (PeproTech). iPSC clones were analyzed by qRT-PCR to quantify *OCT4, NANOG, LIN28, DNMT3B* and *Sendai.* Immunofluorescence analyses were performed to examine expression and localization of OCT4, SOX2, SSEA4 and TRA-1-60. iPSC clones were also characterized using the hPSC Genetic Analysis Kit (Stem Cell Technologies). Cell lines were authenticated by STR analysis and tested for mycoplasma using the MycoAlert$^{TM}$Plus Mycoplasma Detection Kit (Lonza).

**Generation of U-87 MG cells with UBA5 pathogenic variants**

Briefly, 500,000 U87-MG cells (ATCC) were nucleofected (Lonza, 4D-ucleofector™ X-unit) with precomplexed ribonuclear proteins (RNPs) consisting of 100 pmol of chemically modified sgRNA (Synthego), 35 pmol of 3X NLS *Sp*Cas9 protein (St. Jude Protein Production Core), and 50 pmol of each ssODN donor in a small (20 µl) cuvette using solution P3 (Lonza) and program EN-158 according to the manufacturer's recommended protocol.  The *UBA5* p.A371T clones were created using Cas12a RNPs consisting of 160 pmol of crRNA (IDT) and 126 pmol of *As*Cas12a protein (IDT). The nucleofections to create the *UBA5* p.A371T clones were performed as above with the addition of 78 pmol of Alt-R® Cpf1 Electroporation Enhancer (IDT). For clones with multiple modifications, modifications were performed sequentially. Cells were single cell sorted five days post nucleofection at the Flow Cytometry and Cell Sorting Shared Resource (St. Jude). Clones were plated into prewarmed (37°C) media into 96-well plates. Clones were screened for the desired modification via gel electrophoresis and targeted deep sequencing on a Miseq Illumina sequencer as previously described (*96)*. NGS analysis of clones was performed using CRIS.py. (*97)*. Correctly modified clones were identified, expanded, and sequence confirmed. Cell identity was authenticated using the PowerPlex® Fusion System (Promega) performed at the Hartwell Center (St. Jude) and tested negative for mycoplasma by the MycoAlert™Plus Mycoplasma Detection Kit (Lonza). Editing construct sequences and relevant primers are listed in Table S2.

**UBA5 sgRNA design and assembly**

sgRNAs were designed and assembled in the Center for Advanced Genome Engineering at St. Jude. Five sgRNAs were designed to target the 5' UTR of *UBA5* and to have at least 2bp of mismatch between the target site and any other site in the human genome. Additionally, a non-targeting sgRNA that cannot bind in the human genome was used as a negative control. sgRNAs were cloned into the pAW12.lentiguide.GFP (Addgene, 104374) using oligos (IDT) with appropriate overhangs (Table S3). Briefly, the pAW12.lentiguide.GFP backbone was digested with Anza Esp3I (ThermoFisher,), followed by heat inactivation. Sense and antisense oligos were mixed at 1:1 ratio from 100 uM stocks. The oligo mixture was then annealed by heating to 95°C for 10 mins followed by slow cooling to room temperature. Oligos mixtures were phosphorylated using T4 PNK (NEB) according to the manufacturer's protocol and diluted 1:200 in water. The diluted oligo mixture was ligated into the digested pAW12.lentiguide.GFP using T4 Ligase (NEB) at room temperature for 1.25 hr according to the manufacturer's recommended protocol. The ligation mixture was transformed into NEB Turbo *E. coli* cells (NEB) by heat shock and plated on carbenicillin plates. Individual colonies were picked, cultured, mini-prepped, and confirmed by Sanger sequencing using a sequencing primer (5'-ACTGTAAACACAAAGATATTAGTAC-3').

#### Generation of U-87 MG cells with dCas9$^{-VPR}$

U87-MG cells were transduced with lentiviral vector packaged with lenti-EF1a-dCas9-VPR-Puro plasmid (Addgene, 99373) at an MOI of 1 with LentiBOOST (Sirion Biotech). Next day, the growth medium was changed and puromycin selection (2 ug/mL) was included in the growth media for the next two weeks to select for expression of dCas9$^{-VPR}$. After selection, the expression of dCas9$^{-VPR}$ was confirmed by western blot analysis of Cas9 (Millipore). Cells were then maintained in puromycin (1 µg/mL) for all experiments.

#### Long-read sequencing

Targeted long-read sequencing (LRS) on Oxford Nanopore Technologies (ONT) platform was used to validate that the p.R55H and p.A371T variants were correctly edited on separate alleles (in trans) in U-87 MG cells. Targeted LRS using "read-until" function was performed on an ONT GridION using a single R9.4.1 flowcell as described previously (*98*). At least 100Kb of sequence was added to either side of the target region for capture. All data were base-called using Guppy 6.3.2 (ONT). Reads were aligned to GRCh38/hg38 using minimap2 and phased bam files were visualized in IGV.

#### Lentiviral vector production and titration

SJ293TS cells were transfected with pCAG-kGP1-1R, pCAG-VSVG and pCAG4-RTR2 using PEIpro (Polyplus Transfection). The following day, transfected cells were diluted with media containing 12.5 U/mL Benzonase (Sigma). Vector supernatants were collected 48hr post-transfection, clarified by centrifugation at 330G for 5 mins and filtered. Lentiviral vector containing supernatants were adjusted to

300 mM NaCl, 50mM Tris pH 8.0 and loaded onto an Acrodisc Mustang Q membrane (Pall Life Sciences) according to the manufacturer's instructions using an Akta Avant chromatography system (GE Healthcare). After washing the column with 300 mM NaCl, 50 mM Tris pH 8.0, viral particles were eluted from the column using 2 M NaCl, Tris pH 8.0 directly onto a PD10 desalting column (GE Healthcare) according to the manufacturer's instructions. Vector containing flow-through was diluted with an equal volume of X-VIVO 10 media (Lonza) or phosphate buffered saline containing 0.5% (v/v) human serum albumin (Grifols Biologics) to achieve an approximate 50-fold concentration from the starting material, sterile filtered, aliquoted and stored at −80°C. Titration of lentiviral vectors was performed by transducing HOS cells with serially diluted vector preparations in the presence 5-8 mg/mL Polybrene (Sigma). Four days post-transduction, genomic DNA is prepared by lysing cells directly in culture plate (35,000 cells/100 uL lysis buffer), then proteinase K treating each sample prior to digital droplet PCR titration. Lysis/extraction buffer contains 10mM Tris pH 8.0, 2 mM EDTA pH 8.0, 0.2% Triton X-100, 200 μg/ml proteinase K. Vector titers were determined by calculating the ratio between the copies of HIV psi and every two copies of RPP30 via QX200 digital droplet PCR system (Bio-Rad), multiplied by the number of cells transduced and if necessary, multiplied by the dilution factor.

**Lentiviral transduction**

Lentivirus vectors were added to the medium with LentiBOOST (Sirion Biotech) according to manufacturer's instructions and incubated overnight. For U-87 MG cells, SINEUP and sgRNA lentiviral vectors were transduced at an MOI of 10, and the medium was changed 16 hr after transduction. For SINEUP experiments in CO, the medium was changed 24 hr after transduction.

**Western blot analysis**

Whole cell lysates were obtained by lysing frozen cell pellets or live cells in RIPA buffer (Cell Signaling Technology) with protease/phosphatase inhibitors (Cell Signaling Technology). Cell lysates were left on rotation for 15 mins at 4°C. Lysates were clarified by centrifugation at 16,000G for 10 mins at 4°C, and protein concentration was determined using the BCA assay (Pierce, Thermo Fisher Scientific). Clarified lysates were then mixed with SDS sample buffer (Bio-Rad), resolved on SDS-PAGE gels (Bio-Rad), transferred to nitrocellulose membranes, and then blocked in TBS-Tween with 3% (w/v) bovine serum albumin (Sigma). Membranes were incubated with indicated primary antibodies overnight at 4°C and detected by HRP-conjugated secondary antibodies and enhanced chemiluminescence (GE Healthcare). All images shown in the manuscript are representative images of 3-5 independent experiments. Quantification were performed first by normalizing target protein to GAPDH, then normalized to indicated control.

**RNA preparation and qRT-PCR**

RNA was extracted from cultured cells using *Quick*-RNA Miniprep Kit (Zymo Research) according to the manufacturer's instructions. A total of 2 μg of total RNA was converted to cDNA using SuperScript VILO Master Mix (Thermo Fisher Scientific). Quantitative PCR reactions were run in triplicate in a QuantStudio 6 Real-Time PCR system (Thermo Fisher Scientific). Analysis of results was done using the ΔΔCt method and normalized to *ACTB*. Primers are listed in Table S4.

**Immunofluorescence, image acquisition, and image analysis of U-87 MG cells**

Cells were fixed with 4% (*v/v*) paraformaldehyde for 10 mins at room temperature and permeabilized with PBS containing 0.25% (*v/v*) Triton X-100. Cells were blocked in PBS with 0.1% (*v/v*) Triton X-100 and 3% BSA for 1 hr at room temperature. Antibodies were diluted in PBS with 0.1% (*v/v*) Triton X-100 and

1% BSA, with primary antibodies incubated overnight at 4°C, and secondary antibodies incubated for 1 hr at room temperature. Cells were washed 3 times for 10 mins each in PBS containing 0.5% (*v/v*) Tween-20 after incubations. Slides were mounted with Aqua-Mount Mounting Medium (VWR). Fixed cells were imaged using a 63 × 1.4 NA oil objective (Leica) with the LAS X software (Leica) on the Leica TCS SP8 (Leica).

Following image acquisition, separation of the fluorescent channels was performed in ImageJ v1.54f (NIH). Briefly, analysis of ER area was performed by first using Ilastik to identify and classify calnexin signal, phalloidin signal, nuclear signal, and background signal. Then, segmented images along with original images were analyzed in CellProfiler to quantify calnexin area and phalloidin area in each image.

**Immunofluorescence and clearing of whole CO**

CO were fixed with 4% (v/v) paraformaldehyde for overnight at 4°C under rotation. CO were then preserved and delipidated using the Clear+ Passive Clearing Kit (LifeCanvas Technologies) according to manufacturer's instructions. Briefly, after SHIELD formation, CO were delipidated in Delipidation buffer for three days at 37°C under rotation. Next, CO were blocked in PBS with 0.1% (v/v) Triton X-100 and 3% BSA for 6 hr at room temperature. Antibodies were diluted in PBS with 0.1% (v/v) Triton X-100 and 1% BSA, incubated for 3 days at room temperature. CO were washed 3 times for 1 hr each in PBS containing 0.5% (v/v) Tween-20 after incubations. Refractive index matching of CO were performed using EasyIndex (LifeCanvas Technologies) according to manufacturer's instructions. CO were then mounted in 2% low melting point agarose in EasyIndex in glass capillaries (SIGMA Z328510) one day prior to light sheet microscopy and kept at 4°C.

**Light sheet microscopy of cleared CO**

Whole CO were imaged on a Zeiss Z1 light sheet microscope equipped with a 5x detection objective (NA 0.16) and two 5x light sheet forming objectives (NA 0.1). Samples were suspended in EasyIndex (RI=1,52) in LMP agar and first excited simultaneously with 488nm and 639 nm followed by excitation with 561 nm. Band-pass emission filters used were: 488 – BP525-565; 561 – BP575-615; 639 – LP660. Z-stacks were spaced at 5 μm apart, and tiled images were aligned and fused using the FIJI BigStitcher plugin. Analysis of SATB2 and CTIP2 staining density was performed using FIJI macros. Briefly, 2D maxima finding was applied to each slice as a proxy for presence of nuclear staining followed by manual thresholding on the density of the 2D peak localizations. Image resolution was insufficient to accurately count individual cells. The volume of above threshold regions was then normalized to the cellular volume of the organoid as defined by ToPro3 staining. In **Fig. 3C**, maximum-intensity projection of the middle 20 slices was presented.

**Supplemental Tables**

**Table S1.** Marker genes used to annotate cell clusters for cortical organoids.

| Cluster 1 | Cluster 2 | Cluster 3 | Cluster 4 | Cluster 5 | Cluster 6 |
|---|---|---|---|---|---|
| SCGN | GPR22 | PPP1R17 | CXCL14 | HSPA1B | VIM |
| CALB2 | IGFBP5 | NHLH1 | SFRP2 | DNAJB1 | METRN |
| SYNPR | ATP8A2 | EOMES | CCL2 | DUSP8 | HOPX |
| DOCK5 | SLC24A2 | NEUROD4 | LIX1 | OSER1 | CLU |
| DLX5 | SYNDIG1 | SSTR2 | FGFR3 | TAF1D | S100B |
| ST8SIA5 | PDE1A | RASGEF1B | IFI44L | VEGFA | ANXA1 |
| SP9 | CACNA1E | FOXP4 | F3 | NRN1 | GFAP |
| DLX6 | MEF2C | TMEM158 | SLCO1C1 | ANKRD37 | TTYH1 |
| NXPH1 | NPR3 | PLCB4 | ID3 | EIF1 | PTN |
| ERBB4 | OPCML | UNC5D | C1QL1 | WDR33 | ANXA5 |
| SLC32A1 | INHBA | SETD7 | ITGA2 | CLK1 | EMP3 |
| RGN | RPRML | KCNQ3 | CRH | SH3BP5 | MT3 |
| DLX1 | ARPP21 | ELAVL2 | LTBP1 | ATF4 | S100A11 |
| RBP1 | SIAH3 | PTCHD2 | HES1 | UIMC1 | TAGLN2 |
| NRXN3 | OSTN | ENC1 | LRRC10B | NELL2 | LGALS3 |
| FAM222A | SATB2 | IGDCC3 | SLC1A3 | DSEL | MT2A |
| SP8 | KCNQ5 | DPY19L1 | DIPK1C | ZNF581 | HSPB1 |
| DLX2 | FAM49A | EPHA3 | IQGAP2 | F5H423 | SPARCL1 |
| VAX1 | SERPINI1 | HES6 | STOX1 | NEUROD6 | PEA15 |
| SLC6A1 | SCN2A | PRDX1 | C21orf62 | KCNK12 | CRYAB |
| PDE5A | VSTM2B | SORBS2 | HOPX | RBM33 | C1QL1 |
| PDZRN4 | SHISA2 | TBR1 | ZFP36 | CCNB1IP1 | APOE |
| PBX3 | VSNL1 | IGFBPL1 | SOX3 | PRDM8 | S100A16 |
| SLC4A4 | SPINT2 | SLA | NOG | RSL1D1 | ID3 |
| WNT7A | SNCA | CNR1 | EFHD1 | SULT4A1 | TNFRSF12A |
| MAFB | CHST15 | BHLHE22 | ATP1A2 | GOLGA8A | HTRA1 |
| IL1RAPL1 | GABBR2 | NEUROG2 | CRB2 | RLF | RHOC |
| PLS3 | RNF174 | PLXNA4 | DAAM2 | CDC42EP3 | MGST1 |
| C11orf96 | PCLO | NGDN | HEPN1 | MT-RNR1 | PSAT1 |
| GPD1 | NR4A3 | MLLT3 | SOCS3 | RSRC2 | DBI |
| GAD2 | FGF12 | FRMD4B | MLC1 | CREBRF | PON2 |
| STC2 | KIAA1644 | KLHL35 | PLPP3 | NFIL3 | IFITM3 |
| C6orf32 | NPM2 | SCRT2 | PON2 | TXNL1 | TIMP1 |
| GAD1 | GRIN2B | NRN1 | FAM107A | H4C8 | PCLAF |

| | | | | | |
|---|---|---|---|---|---|
| METAP1D | KIT | OCIAD2 | FZD8 | COLEC11 | TYMS |
| ARX | NYAP2 | NR2F1 | GLI3 | SGF29 | TOP2A |
| SMOC1 | KCNK12 | ELAVL4 | TNC | STMN4 | SPARC |
| PDZRN3 | MAL2 | SOX11 | MRC2 | PDK1 | BBOX1 |
| TMEFF2 | HPCA | NEUROD2 | NR2E1 | PLCG2 | RAMP1 |
| KITLG | DAB1 | ZBTB18 | HLA-E | KMT5B | UBE2C |

| **Cluster 7** | **Cluster 8** | **Cluster 9** | **Cluster 10** | **Cluster 11** | **Cluster 12** |
|---|---|---|---|---|---|
| COL1A2 | STMN2 | CCN2 | NEUROD4 | MGP | GDF10 |
| NPY | TUBA1A | CAV1 | EOMES | CYP26A1 | CFAP126 |
| BEST3 | NEUROD6 | DNAJB1 | NEUROG1 | BARHL1 | IFITM1 |
| GSX2 | NEUROD2 | HKDC1 | DLGAP5 | MAB21L1 | COL21A1 |
| HELT | MEF2C | HSPA6 | CENPE | RELN | MGST1 |
| PDGFRA | CNTN1 | BHLHE40 | NUF2 | CBLN1 | RSPO2 |
| GJA1 | OSTN | SPP1 | UBE2C | CPLX3 | LUM |
| UBE2C | NSG2 | SLC16A3 | RRM2 | PAX2 | FRZB |
| CDK6 | EEF1A2 | TNFRSF12A | STXBP6 | ZIC4 | CP |
| CDCA8 | CALM1 | SAT1 | CDCA8 | EBF3 | PIK3C2G |
| TOP2A | TTC9B | CRACDL | MYBL1 | UNCX | PAX3 |
| RASD1 | MAL2 | VEGFA | TPX2 | FSTL4 | IRX3 |
| KNL1 | IGFBP5 | P3H2 | H2AC14 | TMEM71 | C1QTNF3 |
| H3C2 | PDE1A | HSPA1B | CENPF | CADPS2 | CYP26B1 |
| GAD2 | CSRP2 | P4HA1 | NDC80 | MRLN | FIBIN |
| HMGB2 | NELL2 | TPD52L1 | CCNB2 | ADCYAP1 | GSN |
| MKI67 | SNCA | CCN1 | DEPDC1 | NDST3 | RSPH1 |
| DLGAP5 | RTN1 | AK4 | CDC25C | EN2 | DYNLT5 |
| SGO1 | PCSK1N | PAWR | BUB1B | BMP5 | ABCA8 |
| CENPE | SERPINI1 | BNIP3 | TOP2A | ATOH1 | TPPP3 |
| KIF4A | NEFL | SERPINH1 | HMMR | NPTX2 | TPBG |
| CKAP2L | GAP43 | NRG1 | ESCO2 | FGF5 | OLFML1 |
| CENPF | UCHL1 | LRATD2 | ASPM | EBF1 | EFCC1 |
| NUF2 | GPR22 | SPRY2 | CEP55 | MYCT1 | CALB1 |
| SGO2 | NEFM | PLOD2 | KIFC1 | HSD11B2 | ARMC3 |
| PTTG1 | NSG1 | SAMD4A | PCLAF | HS3ST5 | GDPD2 |
| GTSE1 | BEX1 | RASSF8 | GTSE1 | CDH7 | ITIH5 |
| ASPM | CDKN2D | SPRY1 | SPC25 | HOXB2 | ELN |
| PBK | SPINT2 | SLC3A2 | H1-5 | UNC5C | IRX5 |
| KIFC1 | RPRML | CEBPB | AURKA | ESRRG | COL12A1 |
| NDC80 | SMIM43 | MTHFD1L | SGO1 | DMKN | VEGFC |

| | | | | | |
|---|---|---|---|---|---|
| CCNB2 | SNAP25 | CIART | HMGB2 | KCNA1 | CD36 |
| KIF11 | CAMK2B | SLC2A1 | KNL1 | IRX2 | ECM2 |
| TPX2 | JPT1 | RORA | MKI67 | BARHL2 | SPARCL1 |
| SPC24 | LINGO1 | PDK1 | MAD2L1 | AKAIN1 | LRP2 |
| BIRC5 | SYT1 | FAM162A | CENPA | ZIC1 | MORN5 |
| NUSAP1 | PPP2R2B | CD9 | PARPBP | CMTM7 | ANGPT1 |
| ESCO2 | CYRIA | GNG12 | CKAP2L | GRIK2 | ZMYND10 |
| SPC25 | MAPT | EMP3 | ECT2 | NKD1 | CAPSL |
| CDKN3 | VSNL1 | DDIT4 | TTK | PCSK9 | SFRP4 |

**Table S2.** CRISPR-Cas9 editing construct sequences.

| **Name** | **Sequence (5' to 3')** |
|---|---|
| *UBA5* p.A371T reagents | |
| CAGE1072.UBA5.Cas12a.g1 spacer | AAGUACCUUUUUUGGAAUUGU |
| CAGE1072.DS.F | CTACACGACGCTCTTCCGATCTtgaggtttcagaagaggaactga |
| CAGE1072.DS.R | CAGACGTGTGCTCTTCCGATCTtgcatattaacactgcatacctctc |
| CAGE1072.g1.anti.ssODN<br>*Alt-R modifications | *attttatatgatatttacatatgggtaaatcatatttttgaagtacTttCtttggaattgtgtatgTcactgtaattccttcaggtaagtctggaactggacctgaa |
| CAGE1072.g1.BLOCK.anti.ssODN<br>*Alt-R modifications | *attttatatgatatttacatatgggtaaatcatatttttgaagtacTttCtttggaattgtgtatgccactgtaattccttcaggtaagtctggaactggacctgaa |
| *UBA5* fs_p.303X reagents | |
| CAGE1073.UBA5.g1 spacer | GGUACUGUUAGUUUUUACCU |
| CAGE1073.DS.F | CTACACGACGCTCTTCCGATCTtctggatgcagatttctgttttct |
| CAGE1073.DS.R | CAGACGTGTGCTCTTCCGATCTatattcctcctgctgcttcctg |
| CAGE1073.g1.anti.ssODN<br>*Alt-R modifications | *gcttcatggacatagtaggaaaaaaatcctgcattgcattTtatccTAAgtaaaaactaacagtaccaaaatttaacagaaacctagaaa |
| CAGE1073.g1.BLOCK.anti.ssODN<br>*Alt-R modifications | *gcttcatggacatagtaggaaaaaaatcctgcattgcattgtatccTAAgtaaaaactaacagtaccaaaatttaacagaaacctagaaa |
| *UBA5* p.R55H reagents | |
| CAGE1074.UBA5.g1 | CuuuGuuuuAAGCCGCuuGA |
| CAGE1074.UBA5.DS.F | CTACACGACGCTCTTCCGATCTcagcctgtaagcatcctcctc |
| CAGE1074.UBA5.DS.R | CAGACGTGTGCTCTTCCGATCTgctacggcaaaggtacggatt |
| CAGE1074.g1.sense.ssODN<br>*Alt-R modifications | *gactgtaagctttaaaacatatttttctttgttttaagccAcCtCatggcattgaaacgaatgggaattgtaagcgactatga |
| CAGE1074.g1.BLOCK.sense.ssODN<br>*Alt-R modifications | *gactgtaagctttaaaacatatttttctttgttttaagccgcCtCatggcattgaaacgaatgggaattgtaagcgactatga |

**Table S3.** *UBA5* sgRNA sequences.

| Oligo | Oligo sequence (5' to 3') lowercase overhang; uppercase target |
|---|---|
| CAGE2339.UBA5.g1.F | caccg TCTTAGATACCCGGTTAGCA |
| CAGE2339.UBA5.g1.R | aaac TGCTAACCGGGTATCTAAGA c |
| CAGE2339.UBA5.g2.F | caccg GGTTAGCAAGGCAACATCGC |
| CAGE2339.UBA5.g2.R | aaac GCGATGTTGCCTTGCTAACC c |
| CAGE2339.UBA5.g3.F | caccg GCCCTGGACACTTATCACCC |
| CAGE2339.UBA5.g3.R | aaac GGGTGATAAGTGTCCAGGGC c |
| CAGE2339.UBA5.g4.F | caccg CACCGGGGTGATAAGTGTCC |
| CAGE2339.UBA5.g4.R | aaac GGACACTTATCACCCCGGTG c |
| CAGE2339.UBA5.g5.F | caccg GCGATGTTGCCTTGCTAACC |
| CAGE2339.UBA5.g5.R | aaac GGTTAGCAAGGCAACATCGC c |
| CAGE464.NT.g10.F | caccg CCTTAACGGCAATCGCGCGT |
| CAGE464.NT.g10.R | aaac ACGCGCGATTGCCGTTAAGG c |

**Table S4.** qRT-PCR Primers used in this study.

| Primer Name | Sequence |
|---|---|
| ACTB-F | ACCATGGATGATGATATCGC |
| ACTB-R | TCATTGTAGAAGGTGTGGTG |
| UFC1 F | TTTGGACTAGCTCATCTCATGG |
| UFC1 R | GAATCAGATCAGGGATTTCCAC |
| UFM1 F | GGCTGCCGTACAAAGTACTCA |
| UFM1 R | TTCCATCATTGGTAATAATTGCAC |
| SOX2-F | TTCACATGTCCCAGCACTACCAGA |
| SOX2-R | TCACATGTGTGAGAGGGGCAGTGTGC |
| MAP2-F | CCACCTGAGATTAAGGATCA |
| MAP2-R | GGCTTACTTTGCTTCTCTGA |
| Foxg1-F | GTATGTGGTCACTAACAGGTC |
| Foxg1-R | ACCACAGTATCACAATCAAG |
| Pax6-F | CTGAGGAATCAGAGAAGACAGGC |
| Pax6-R | ATGGAGCCAGATGTGAAGGAGG |
| Oct4-F | CCCCAGGGCCCCATTTTGGTACC |
| Oct4-R | ACCTCAGTTTGAATGCATGGGAGAGC |
| LIN28-F | AGCCATATGGTAGCCTCATGTCCGC |
| LIN28-R | TCAATTCTGTGCCTCCGGGAGCAGGGTAGG |
| Nanog-F | AGATGCCTCACACGGAGACT |
| Nanog-R | TTGGGACTGGTGGAAGAATC |
| DNMT3B-F | ATAAGTCGAAGGTGCGTCGT |
| DNMT3B-R | GGCAACATCTGAAGCCATTT |
| Sendai-F | GGATCACTAGGTGATATCGAGC |
| Sendai-R | ACCAGACAAGAGTTTAAGAGTATGTATC |
| Emx2-F | CACGGAAACTCAGGTAAAAG |
| Emx2-R | CGGTTAATATGGTGCGTC |

| | |
|---|---|
| Emx1-F | GCCTTCGAGAAGAACCACTACG |
| Emx1-R | CGGTTCTGGAACCACACCTTCA |
| Fezf2-F | CGCTCAACACGCATATCC |
| Fezf2-R | GCTTGTGGTTCTTGTAGTTCC |
| Ctip2-F | ACATGAAAAAGTGGCACGGC |
| Ctip2-R | GAGGCAAGTCAGGTCAGCAT |
| Satb2-F | CAAGAGTGGCATTCAACCGCAC |
| Satb2-R | ATCTCGCTCCACTTCTGGCAGA |
| UBA5 F | TTGCCCAGGAGAGGAGTCTG |
| UBA5 R | CATCTTCTCGATGCGGACCC |
| DCX F | TATGCGCCGAAGCAAGTCTCCA |
| DCX R | CATCCAAGGACAGAGGCAGGTA |
| Tuj1 F | TCAGCGTCTACTACAACGAGGC |
| Tuj1 R | GCCTGAAGAGATGTCCAAAGGC |
| GAT1 F | AACACAGACCGCTGCTTCTCCA |
| GAT1 R | AGCGGATCTGACCTGGCTTATC |
| GAD2 F | GCCAACTCTGTGACGTGGAATC |
| GAD2 R | GCTGAAAGAGGTAGGAGGCATG |
| GAD1 F | TGTCCAGGAAGCACCGCCATAA |
| GAD1 R | TCCTTGACGAGAATGGCAGAGC |
| GABRA1 F | CACAAGTCTCCTTCTGGCTCAAC |
| GABRA1 R | GGAGTTTCTGGCACTGATGCTC |
| SLC32A1 F | CTGGAACGTGACCAACGCCATC |
| SLC32A1 R | TCATTCTCCTCGTACAGGCACG |
| GRIN1 F | CCAGTCAAGAAGGTGATCTGCAC |
| GRIN1 R | TTCATGGTCCGTGCCAGCTTGA |
| GRIN2B F | TTCCACTGGCTATGGCATTGCC |
| GRIN2B R | GACAAATGCCAGTGAGCCAGAG |
| VGLUT2 F | GAGAGGAGTAGACTGGCAACCA |
| VGLUT2 R | CTGAAGACCAGCCAGTGTACTG |
| VGLUT1 F | GCAAGTACATCGAGGACGCCAT |
| VGLUT1 R | GCCACGATGATGGCATAGACTG |
| XBP1 F1 | TGGCCGGGTCTGCTGAGTCCG |
| XBP1 R1 | ATCCATGGGGAGATGTTCTGG |
| us XBP1 F1 | CAGCACTCAGACTACGTGCA |
| us XBP1 R1 | ATCCATGGGGAGATGTTCTGG |
| s XBP1 F1 | CTGAGTCCGAATCAGGTGCAG |
| s XBP1 R1 | ATCCATGGGGAGATGTTCTGG |

**Table S5.** Antibodies used in this study.

| Antibody | Vendor | Catalog Number |
|---|---|---|
| ATF6 | Abcam | ab122897 |
| ATF6 | Cell Signaling Technology | 65880 |

| | | |
|---|---|---|
| Beclin | Cell Signaling Technology | 3495 |
| BiP | Cell Signaling Technology | 3183 |
| Calnexin | Thermo Fisher Scientific | MA3-027 |
| CHOP | Abcam | ab11419 |
| cleaved PARP | Cell Signaling Technology | 5625 |
| CTIP2 | Abcam | ab18465 |
| EIF2α | Cell Signaling Technology | 5324 |
| GAPDH | Cell Signaling Technology | 2118 |
| Grp94 | Cell Signaling Technology | 2029 |
| HERPUD1 | Cell Signaling Technology | 26730 |
| IRE1α | Cell Signaling Technology | 3294 |
| PARP | Cell Signaling Technology | 9542 |
| PERK | Cell Signaling Technology | 5683 |
| Phalloidin | Thermo Fisher Scientific | A22287 |
| p-IREα | Novus | NB100-2323 |
| p-EIF2α | Cell Signaling Technology | 3398 |
| p-PERK | Thermo Fisher Scientific | PA540294 |
| UBA5 | Abcam | ab177478 |
| UFC1 | Abcam | ab189252 |
| UFBP1 | Abcam | ab251704 |
| UFL1 | Abcam | ab22616 |
| UFM1 | Abcam | ab109305 |
| SATB2 | Cell Signaling Technology | 39229 |
| SATB2 | Abcam | ab34735 |
| GAD2 | Cell Signaling Technology | 5843 |
| GAT1 | Cell Signaling Technology | 37342 |
| PSD95 | Cell Signaling Technology | 3450 |
| TUJ1 | Cell Signaling Technology | 5568 |
| OCT4 | Thermo Fisher Scientific | A24867 |
| SSEA4 | Thermo Fisher Scientific | A24866 |
| SOX2 | Thermo Fisher Scientific | A24759 |
| TRA-1-60 | Thermo Fisher Scientific | A24868 |
| Anti-Mouse IgG HRP-linked | Cytiva | NA931V |
| Anti-Rabbit IgG HRP-linked (from donkey) | Cytiva | NA934V |
| Alexa Fluor 488 | Thermo Fisher Scientific | A32723 and A32731 |
| Alexa Fluor 546 | Thermo Fisher Scientific | A11030 and A11035 |
| Alexa Fluor 647 | Thermo Fisher Scientific | A32733 and A32728 |
| DAPI | Millipore Sigma | D4592 |

| | | |
|---|---|---|
| TO-PRO-3 | Thermo Fisher Scientific | T3605 |
